# Origin and segregation of the human germline

**DOI:** 10.1101/2022.07.06.498671

**Authors:** Aracely Castillo-Venzor, Christopher A. Penfold, Michael D. Morgan, Walfred W. C. Tang, Toshihiro Kobayashi, Frederick C. K. Wong, Sophie Bergmann, Erin Slatery, Thorsten E. Boroviak, John C. Marioni, M. Azim Surani

## Abstract

Human germline-soma segregation occurs during weeks 2-3 in gastrulating embryos. While direct studies are hindered, here we investigate the dynamics of human primordial germ cell (PGCs) specification using in vitro models with temporally resolved single-cell transcriptomics and in-depth characterisation to in vivo datasets from human and non-human primates, including a 3D marmoset reference atlas. We elucidate the molecular signature for the transient gain of competence for germ cell fate during peri-implantation epiblast development. Further, we show that both the PGCs and amnion arise from transcriptionally similar TFAP2A positive progenitors at the posterior end of the embryo. Notably, genetic loss of function experiments show that TFAP2A is crucial for initiating the PGC fate without detectably affecting the amnion, and its subsequently replaced by TFAP2C as an essential component of the genetic network for PGC fate. Accordingly, amniotic cells continue to emerge from the progenitors in the posterior epiblast, but importantly, this is also a source of nascent PGCs.

## Introduction

Human primordial germ cells (PGCs) are among the first lineages to emerge in the developing gastrulating peri-implantation embryo at weeks (Wks) 2-3, eventually developing into sperm or eggs. The parental gametes generate the totipotent state at fertilisation and transmit genetic and epigenetic information necessary for development.

The specification of PGCs is linked with the initiation of the unique germ cell transcriptomic and epigenetic program. Aberrant specification and development of germ cells can lead to sterility, germ-cell-derived cancers, and other human diseases with long term consequences across generations. Ethical and technical reasons restrict direct studies on nascent human PGCs, necessitating in vitro models, which are, however, experimentally tractable for mechanistic insights (Hirate *et al*., 2013; Irie *et al*., 2015; Sasaki *et al*., 2015; Tang *et al*., 2016; Kobayashi *et al*., 2017; Irie, Sybirna and Surani, 2018). Due to the in vitro nature of these models, comprehensive comparisons with rare human embryos and animal proxies, including in vivo development of non-human primates such as marmosets can be significantly informative for germline biology.

The induction of PGC-competent cells from cultured pluripotent stem cells (PSCs) is possible using self-renewing or transient pre-mesendoderm (PreME) populations (Irie *et al*., 2015; Sasaki *et al*., 2015; Kobayashi *et al*., 2017). The timing and regulation of the transient state of competence for PGC-fate in human embryos are not yet fully defined but likely determine the number of founder PGCs in vivo. If aggregated into 3D embryoid bodies, these competent cells give rise to 10 - 40% PGC-like cells (PGCLCs) in response to BMP and other cytokines (Irie *et al*., 2015; Sasaki *et al*., 2015; Kobayashi *et al*., 2017). The remaining cells adopt somatic fates, but their relationship with the emerging PGCLCs remains unclear (Irie *et al*., 2015; Sasaki *et al*., 2015; Kobayashi *et al*., 2017). Defining the characteristics of the somatic lineages in embryoid bodies may help identify soma-PGC interactions and reveal the context of how PGCs form in experimental models concerning the lineages in the embryo.

In vitro models identified *SOX17, PRDM1*, and *TFAP2C* as the core regulators of human PGC fate (Irie *et al*., 2015; Kobayashi *et al*., 2017; Kojima *et al*., 2017; Tang *et al*., 2022). This tripartite network for PGC fate has also been observed in vivo in other species that develop as a bilaminar disc, including cynomolgus, marmoset, rabbit, and pig (Sasaki *et al*., 2016; Sybirna, Wong and Surani, 2019; Alberio, Kobayashi and Surani, 2021; Kobayashi *et al*., 2021; Zhu *et al*., 2021; Bergmann *et al*., 2022).

On the other hand, *Sox17* is not a critical regulator of PGC specification in rodents where the embryos develop as egg cylinders (Kanai-Azuma *et al*., 2002). Notably, when *SOX17* is the critical regulator for PGC specification as in humans and non-human primates, there is concomitant repression of SOX2, but not in mice, where *Sox2* has a crucial role in PGC development (Campolo *et al*., 2013).

The site of human PGC specification remains unclear. In cynomolgus and marmosets, PGCs are first observed in the amnion prior to gastrulation (Sasaki *et al*., 2016; Bergmann *et al*., 2022). At later stages, PGCs are detected in the posterior epiblast, with the possibility of a dual origin (Sasaki *et al*., 2016; Kobayashi *et al*., 2017; Kobayashi and Surani, 2018). Note that in humans and non-human primates, the nascent amnion is among the first lineages to form from the epiblast (Bergmann *et al*., no date; Xiang *et al*., 2019). In some non-primate embryos, including bilaminar disc forming species such as rabbit and pig, PGC specification precedes amnion development (Alberio, Kobayashi and Surani, 2021; Kobayashi *et al*., 2021; Zhu *et al*., 2021). In the pig, at least, PGCs arise from pre-primitive streak (PS) and early-PS stage competent epiblast(Kobayashi *et al*., 2017), and in a rare Wk3 (Carnegie stage 7) human embryo, PGCs are associated with the primitive streak (Tyser *et al*., 2021).Here we used our PSC-based model for PGC specification (Irie *et al*., 2015; Kobayashi *et al*., 2017) in conjunction with highly resolved single-cell transcriptome sequencing and integrative analysis with existing human and primate datasets to document the nature of the somatic components of the models and provide the context for PGCLC specification. Notably, we identified *TFAP2A*, considered an amnion marker (Shao, Taniguchi, Gurdziel, *et al*., 2017; Shao, Taniguchi, Townshend, *et al*., 2017), as an essential and thus far the earliest regulator of PGC fate. Loss of TFAP2A leads to an almost complete abrogation of PGCLCs, in favour of a population of cells displaying SOX2 expression but no significant effect on somatic lineages. The observations also provide insights into the likely origin of human PGCs.

## Results

### A highly resolved transcriptional characterisation of PGC specification in embryoid bodies

Human pluripotent stem cells (PSCs) in a primed state represent non-gastrulating postimplantation epiblast cells (Yu *et al*., 2021) with a low competence for PGCLC fate (<5%) (Irie *et al*., 2015). PSCs can, however, acquire competence for PGC fate as self-renewing populations in media containing four inhibitors (henceforth called 4i conditions) (Gafni *et al*., 2013; Irie *et al*., 2015). Secondly, in response to WNT and Activin signalling, PSC can transiently acquire competence for PGC-fate at 12h, known as pre-mesendodermal cells (henceforth called Pre-ME) (Kobayashi *et al*., 2017). Pre-ME progress to mesendoderm (ME) fate at 24h when they lose competence for PGCLC specification and instead gain competence for definitive endoderm (DE; 60-80%) and mesoderm fates (Fig. 1a).

**Figure 1:**
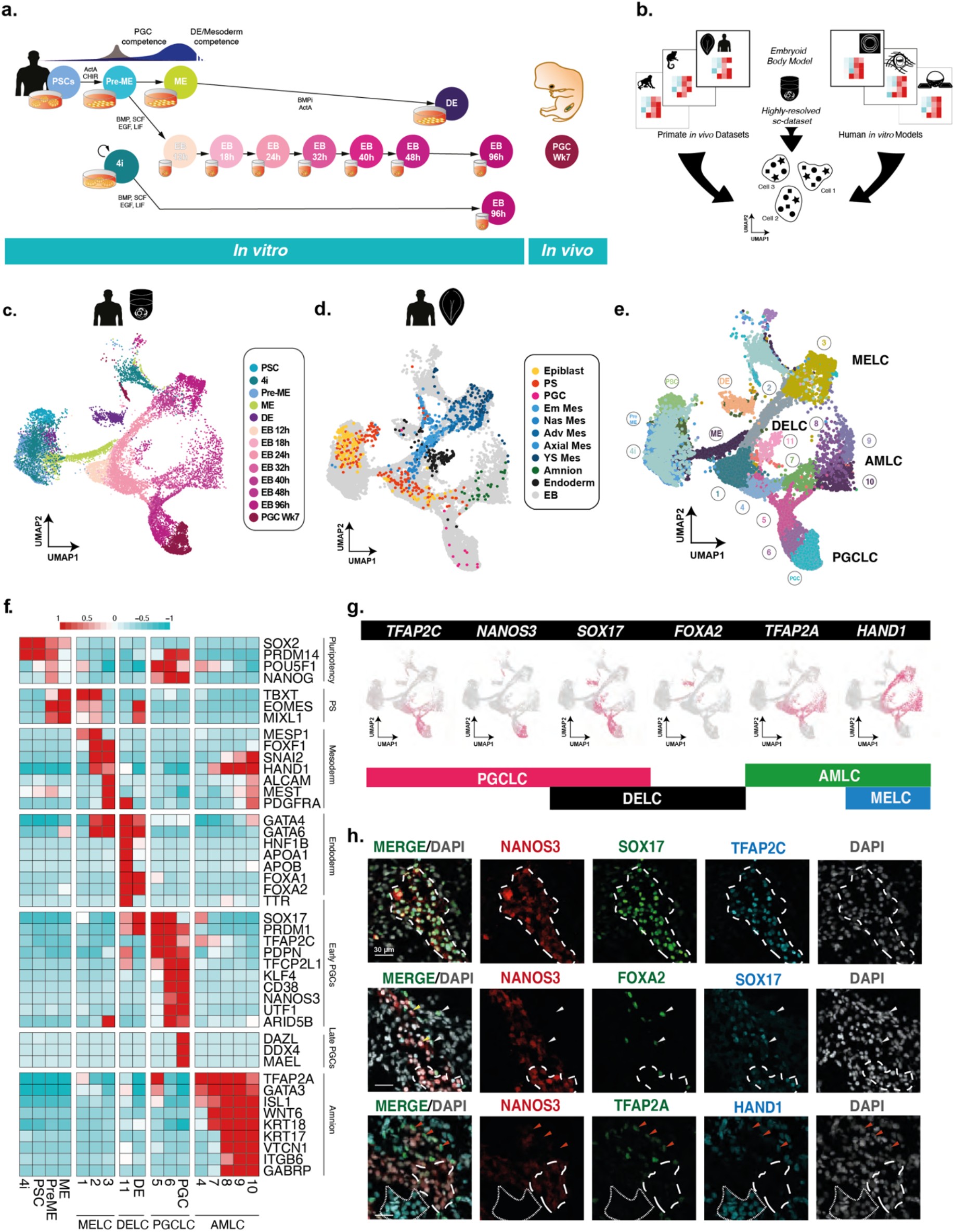
A highly resolved roadmap of PGC development and gastrulation. (a) Experimental design for highly resolved RNA-sequencing (10X) representing an established PGCLC model, PGCLC-competent populations *in vivo* and *in vitro* reference data. PSC; pluripotent stem cells, PreME; pre-mesendoderm (transient PGCLC-competent cells), ME; mesendoderm, 4i; four-inhibitor; self-renewing PGCLC-competent cells, DE; definitive endoderm, PGC; week 7 human gonadal PGCs, EB; embryoid body. (b) Integration of our data with other human *in vitro* models and primate gastrulation datasets used to generate a roadmap of PGC and early human development. (c) Integrated data representation as a UMAP projection with samples highlighted by collection time and sample type. (d) Integrated representation of the aligned human CS7 gastrula data, highlighted by cell type. (e) Louvain clustering of the integrated dataset identified 30 clusters. (f) Heatmaps of pseudobulk expression for key markers showing that the embryoid body diversifies into mesoderm-like cells (MELC), definitive endoderm-like cells (DELC), primordial germ cell-like cells (PGCLC), and amnion like cells (AMLC). (g) Combination of key expression markers with TFAP2C, NANOS3, SOX17 representing PGCLCs; SOX17 and FOXA2 endoderm fate, and TFAP2A and HAND1 to distinguish mesoderm and amnion fates, respectively. (h) Immunofluorescence of d4 EBs confirms PGCLCs, MELCs, AMLCs, and DELCs.

The efficiency of PGCLC induction ranges from ∼10-40% of cells in the embryoid body (EB), depending on the cell line (Chen *et al*., 2017); the remaining non-PGCLCs cells acquire somatic fates. Using our in vitro model (Kobayashi *et al*., 2017), we elucidate the transcriptional dynamics as the Pre-ME cells undergo specification to PGCLCs in response to BMP. To discern the changes in transcriptional states, we analysed the embryoid body at the resolution of single cells using 10X Genomics single-cell RNA-sequencing. We sampled EBs over a highly-resolve time series between 12h–96h post-induction with additional comparative samples of conventional PSCs, PGC-competent populations (4i and PreME), DE and ME populations, and Wk7 human gonadal PGCs for in vivo reference (Fig. 1a).

We first sought to establish the identity of detectable lineages using droplet single cell RNA sequencing in the embryoid bodies, which fell into 15 main clusters (Supplementary Fig. 1a). Pseudo-bulk correlation showed a high degree of correlation between these clusters and amnion-like cells (AMLC), primordial germ-cell like cells (PGCLCs), or mesoderm-like cells (MELCs) (Zheng *et al*., 2019) (Supplementary Fig. 1b). Pseudo-bulk comparison with a human in vitro embryo culture (Xiang *et al*., 2019) and the in vivo CS7 human gastrula (Tyser *et al*., 2021) corroborates these observations, showing a higher degree of correlation between EBs and embryonic disc or amnion but a substantially reduced correlation with other extraembryonic-tissues and pre-implantation lineages (Supplementary Fig. 1c), and a comparatively low correlation with human syncytiotrophoblast (SCT) and extravillous trophoblast (EVT) (Vento-Tormo *et al*., 2018). Together these results suggest that in response to BMP, EBs progress to lineages of the peri-gastrulation embryo but not the extraembryonic tissues except for the amnion.

We aligned our data to a comprehensive range of existing embryonic datasets to refine cell annotations and create a human primate gastrulation and PGC atlas (Fig. 1b). We included embryonic and amniotic lineages from human and cynomolgus in vitro cultured embryos (Ma *et al*., 2019; Xiang *et al*., 2019; Zhou *et al*., 2019), in vivo human and marmoset gastrula (Bergmann *et al*., no date; Tyser *et al*., 2021), and human gonadal primordial germ cells (Guo *et al*., 2015; Li *et al*., 2017). We also include three in vitro models of human PGCLC induction based on the microfluidic amnion model (Zheng *et al*., 2019), micropatterned gastruloids (Minn *et al*., 2020) and embryoid bodies from two other cell lines(Chen *et al*., 2019) (Supplementary table 1). We show a representation of our aligned dataset as a 2D UMAP projection in Fig. 1c, with cells coloured by sampling time. For comparison, we also provide aligned samples from the human gastrula dataset (Fig. 1d), with the remaining datasets shown in Supplementary Fig. 1d-1j. Clustering across all the datasets grouped cells into approximately 30 clusters, with the key clusters visualised for our data in Fig. 1e. Initial assessment suggests a low number of doublets throughout (Supplementary Fig. 1k).

A heatmap of gene expression of relevant lineage markers confirms the presence of a primitive streak-like population (cluster 1), as well as mesoderm-like cells (MELCs) (cluster 2-3), definitive endoderm-like cells (DELCs) (cluster 11), amnion-like cells (AMLCs) (cluster 4, 7-10), and PGC-like cells (PGCLCs) (cluster 5-6) within the EBs (Fig. 1f). We show key differentially expressed transcription factors during the formation of individual cell types in Supplementary Fig. 2. To visualise expression heterogeneity, we depict gene expression of six key lineage markers that, in combination, can be used to identify the cell fates in the EB (Fig. 1g); these findings were also confirmed at the protein level by immunofluorescence (IF) staining (Fig. 1h). Notably, our detailed integrated roadmap and characterisation show that at early stages, embryoid bodies contain subpopulations with molecular signatures similar to the PS, with cells at later time points showing transcriptional profiles associated with embryonic somatic fates (mesoderm and endoderm), PGCLCs, and amnion.

### Detection of PGC competent population

Currently, there is no clear indication of what constitutes a PGC-competent population. We investigate how the precursor PreME cells gain competence for PGC-fate to address this. We also analysed the PGC-competent 4i cells against the non-competent populations (PSCs and ME) (Tang *et al*., 2022). Comparisons with existing datasets suggested that our PSCs are transcriptionally similar to other PSCs(Chen *et al*., 2019; Zheng *et al*., 2019; Minn *et al*., 2020) (Supplementary Fig. 1d-f) and align with a subset of cells from in vitro cultured human embryos labelled as EmDisc (Xiang *et al*., 2019; Zhou *et al*., 2019) (Supplementary Fig. 1g-h) and in vivo postimplantation epiblast (Tyser *et al*., 2021) (Fig. 2-b). Conversely, PreME cells cluster with pluripotent embryonic disc sample (Xiang *et al*., 2019; Tyser *et al*., 2021) and cells labelled as epiblast and primitive streak and mesoderm in a human CS7 gastrula (Tyser *et al*., 2021) (Fig. 2c).

**Figure 2:**
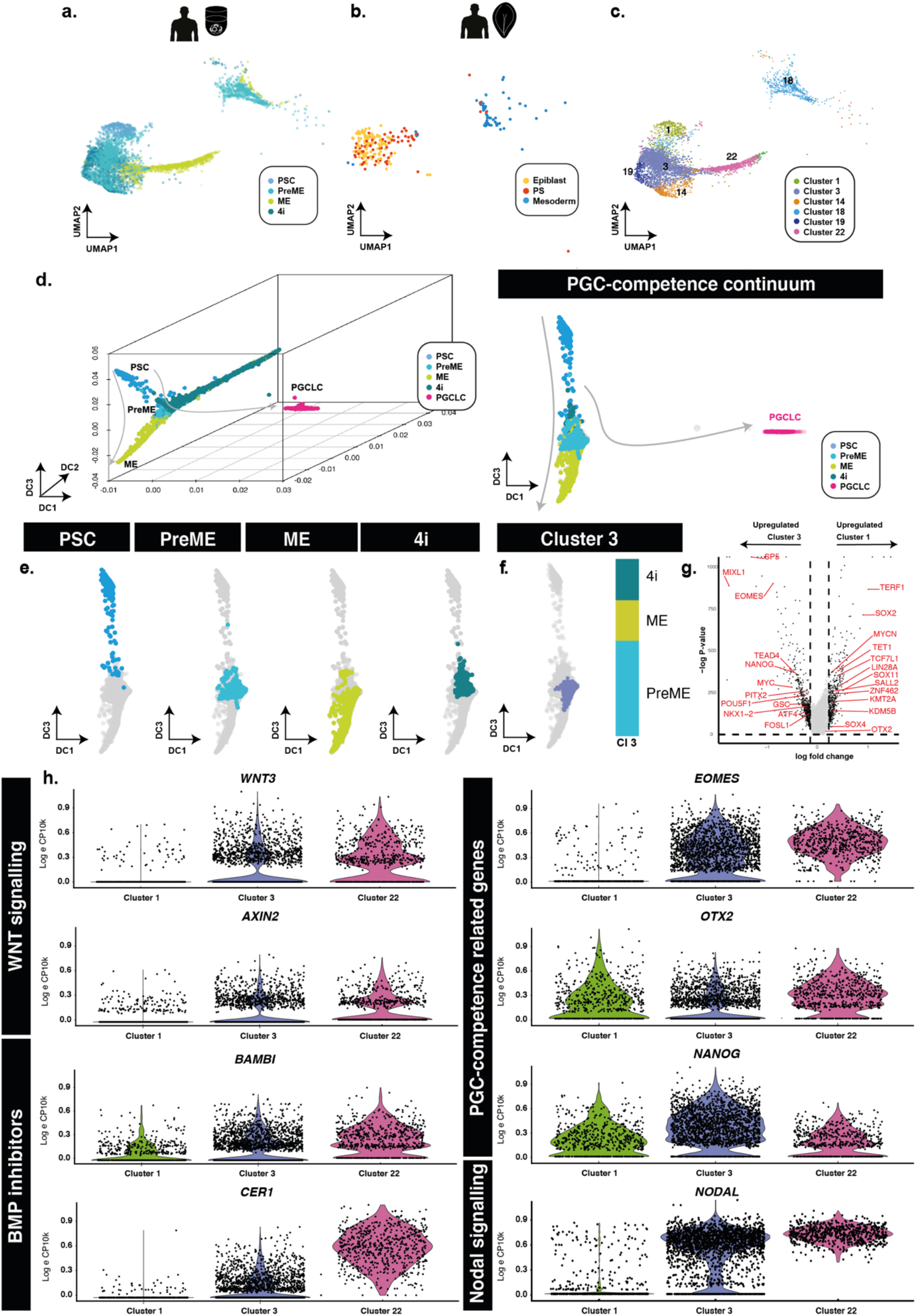
PGCLC competent populations form a continuum of states. (a) Aligned UMAP representations of pluripotent and PGCLC-competent populations, alongside (b), human *in vivo* samples shows that PSCs align best to pluripotent epiblast cells while competent (PreME and 4i) align to both epiblast-like and primitive-streak-like populations. (c) Sub clustering of competent and non-competent cells identified six main populations; a PSC-dominated *cluster 1, a mesendoderm-dominated cluster 22, and competence-dominant clusters 3, 14, 19. (d) Diffusion map representations shows samples lie along a continuum of overlapping states. (e-f) The fractional makeup of competence-dominated subclusters 3 showed almost equal contribution from 4i and PreME cells. (g) Differential expression of competence-dominated subcluster 3 versus PSC-dominant, non-competent subcluster 1 identifies putative regulators of competence. (h) Violin plots of putative competence related genes and markers for WNT and BMP signalling reveal heterogeneous signalling response

Since the interpretation of distances in UMAP representations remains difficult (Chari, Banerjee and Pachter, 2021), we also chose to visualise cells using diffusion maps (DM) to gauge the behaviour of these precursor populations compared to non-competent PSCs, ME populations, and terminal stage PGCLCs (Fig. 2d). These populations exist as a continuum of transcriptional states extending from PSCs to ME, and diffusion components 2 (DC2) and DC3 with PGCLCs extending out along DC1 (Fig. 2e).

Visualisation of the fraction of cell types in each subcluster identified a PSC-dominant (subcluster 1) and a ME-dominant subpopulation (subcluster 22), with three other subpopulations (subclusters 3, 14, and 19) comprised primarily of PGC-competent populations (4i and PreME) (Fig. 2f, Supplementary Fig. 3a). Pairwise differential expression analyses of PreME cells in (competent) subcluster 3 and PSCs in (non-competent) subcluster 1 identified several likely regulators of competence, including *EOMES*, which has an identified role in PGC-competence (Chen *et al*., 2017; Kojima *et al*., 2021), and mesodermal markers *SP5* and *MIXL1* (Fig. 2g). Additional pairwise comparison of the other competent-enriched subpopulations, e.g., subclusters 14 and 19, against cluster 1 identified similar markers, including *OTX2, SOX11, TERF1, TCF7L1, SALL2, LIN28A* and *TET1* (Supplementary Fig. 3b-f). Comparison of competent clusters against cluster 22 showed further upregulation of mesoderm related genes, *MIXL1, GATA6, GSC, MESP1, ZIC2, EOMES* in ME-dominated cluster and concomitant reduction of pluripotency factor expression (*SOX2, SOX3, NANOG*) and MYC in PGC-competent cluster (subcluster 3, 14, 19).

Since competent subclusters 3, 14 and 19 showed similar marker expression (Supplementary Fig. 3b-f), we focused on subcluster 3 for simplicity (Fig. 2f). The signalling dynamics of competence gain and loss were examined by expressing key genes between clusters representing PGC-competent and non-competent cells (Fig. 2f). We observed a progressive activation of NODAL and WNT signalling together with the expression of BMP inhibitors, with the highest levels shown in the ME-dominated subcluster (subcluster 22). BMP is the inductive signal for PGC-fate, and accordingly, the PGC-competent subcluster (subcluster 3) shows a reduced expression of BMP inhibitors *CER1* (p < 2.17e^-19^) compared to the non-competent subcluster 22. Furthermore, we observed upregulation of *NANOG* (p < 9.26e^-48^/1.62e^-6^) and concomitant downregulation of *OTX2* (p < 6.48e^-10^/0.045) in subcluster 3 compared to non-competent clusters 1 and 22 respectively. Notably, *OTX2* negatively regulates PGCLC competence in mice (Zhang *et al*., 2018) and we recently found that *OTX2* has a similar function in the human germline (Tang *et al*., 2022).

Together these analyses have identified molecular signatures that may underlie the transition from primed pluripotency to a competent state for PGC fate.

### Specification of PGCLCs in EBs represents a primitive-streak-like stage

Based on the expression of marker genes, EBs first transition through a primitive-streak-like stage before diversifying into mesoderm-like (MELC), definitive-endoderm-like (DELC), and primordial germ cell-like states, with the additional formation of amnion-like cells but with a notable lack of neural ectoderm populations (Supplementary Fig. 2e). Strikingly, these are lineages expected to arise at the posterior region of the developing embryo around the time of gastrulation.

To test this hypothesis further, we sought to map cells found in vitro to existing spatial transcriptomics datasets. Although spatially resolved human gastruloid datasets exist (Moris *et al*., 2020), these models capture the onset of somitogenesis (CS9) and are therefore more developmentally advanced than our model, which aligns well with data from CS5-7 embryos relevant to the emergence of PGCLCs. In this regard, we note recent comprehensive spatially resolved transcriptional datasets of marmoset embryos at CS5 and CS6 (Bergmann *et al*., 2022), where the peri-implantation development strongly resembles that of human embryo development at the morphological and transcriptional level (Bergmann *et al*., 2022), including conserved expression of *SOX17, PRDM1, TFAP2C* and *NANOS3* in PGCs. Notwithstanding the differences in human and marmoset development timing, archival embryo collections allow consistent staging between species based upon Carnegie staging (Strachan, Lindsay and Wilson, 1997; O’Rahilly and Müller, 2010).

To evaluate possible anterior-posterior bias, we mapped cells from our in vitro model to an existing 3D spatially resolved depiction of a CS6 marmoset embryo in which laser capture microdissection was used to generate a 3D spatially resolved transcriptome (see Materials and Methods). Together, they capture the critical cell types for comparison (Fig. 3a) with gene expression patterns of critical markers shown in Fig. 3b. We found that pluripotent stem cell populations mapped best to the anterior compartment (Fig. 3c), in agreement with earlier studies (Tyser *et al*., 2021; Bergmann *et al*., 2022), although we could not rule out that these cells might have a better mapping to earlier stages, e.g., CS4 bilaminar disc embryo since no data for this stage is available. We found that the PreME population shifted towards the posterior end of the embryo, with amnion-like cells primarily mapping to the posterior amnion (Fig. 3c). The basal cluster, which represents the 12h embryoid body mapped to the posterior end of the embryonic disc to a region expression *TBXT* and other primitive streak markers. Other cell lineages, including PGCLCs, showed an even stronger bias to the posterior end of the embryo, with PGCLC mapping to a distinct *NANOS3*-expressing region between the posterior-most embryonic disc and amnion (Fig. 3a). Together, these results provide further evidence that our model represents the development of the posterior end of the embryo during gastrulation and suggests ongoing specification of both amnion and primordial germ cells.

**Figure 3:**
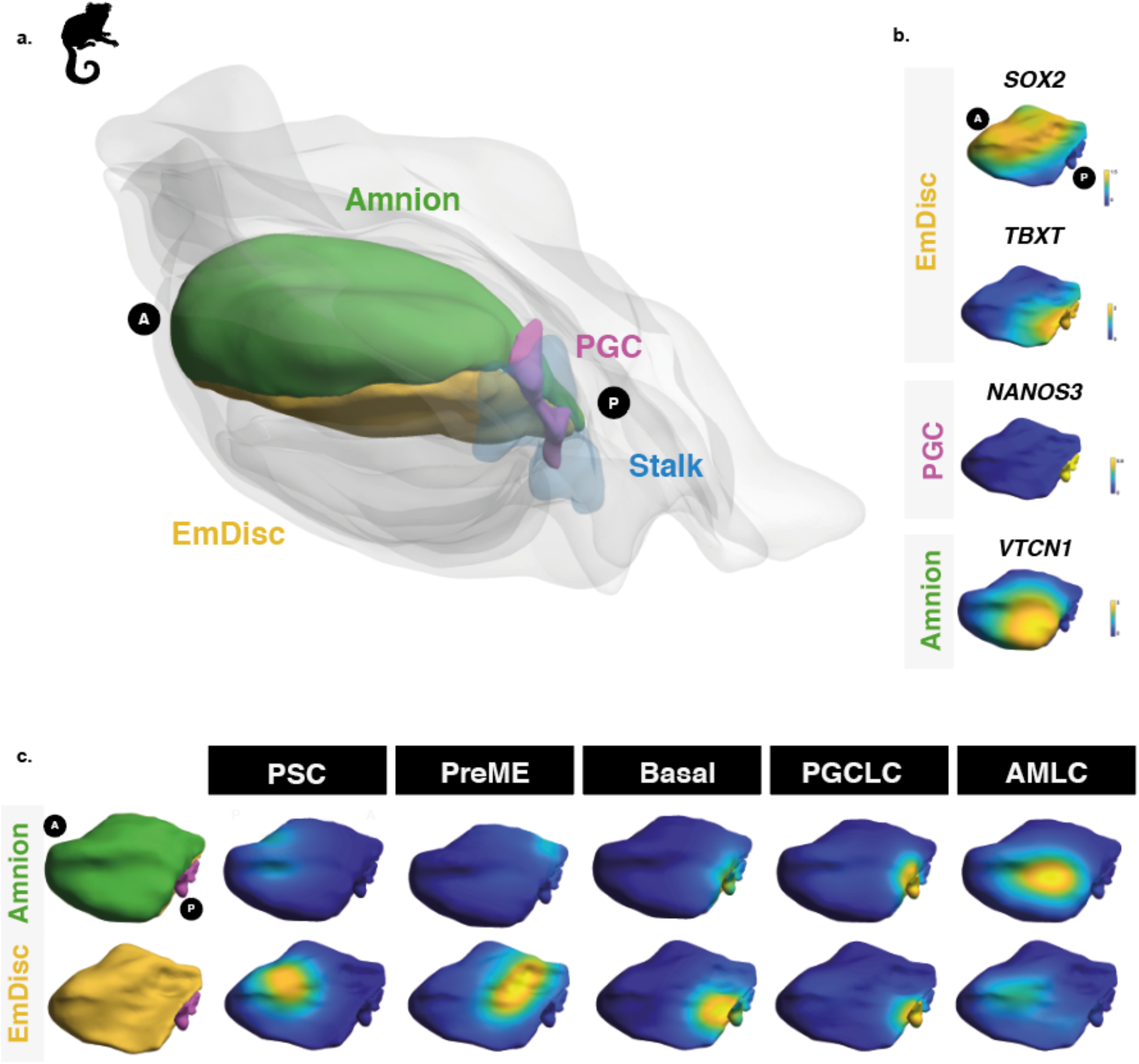
Spatial mapping of embryoid body models to gastrulating marmoset embryos reveals posterior bias. (a) Spatially resolved marmoset embryos at CS6 with the embryonic disc in yellow, amnion in green, PGCs in pink, and stalk in blue. Extraembryonic tissues are shown in grey. (b) Expression analysis in the marmoset embryo shows the anterior embryonic disc is SOX2 and the posterior, T positive, respectively. Specified PGCs with NANOS3 expression, amnion with partial VTCN1 expression. (c) Mapping *in vitro* cells shows PSCs map best to the anterior embryonic disc. Competent populations show a distinct posterior bias, with PGCLCs showing strong localisation to posterior-most PGC region and AMLCs mapping to the amnion.

### Highly resolved time series reveal dynamics of cell trajectories

Having established the identity and spatial correspondence of key lineages, we next investigated the dynamics of individual cell fate decisions within the EB. We performed a label transfer from the human CS7 gastrula dataset (Tyser *et al*., 2021) to our data and separated EBs by collection time to visualise the emergence of cell types (Fig. 4a). Twelve hours after inductive BMP cues, cells aligned primarily to the primitive streak (PS) with a limited pool of epiblast-like cells. Primitive streak-like cells (PSLCs) persisted in limited numbers until ∼24-32h, with sustained expression of *NODAL* (Supplementary Fig. 4). Nascent and emergent mesoderm-like cells (denoted nMELC, eMELC) appeared as early as 12h, becoming more pronounced by 18h, with these lineages roughly corresponding to cluster 2. The earliest PGCLCs arose around the 18h mark, with amnion-like cells and definitive endoderm-like cells around 24h.

**Figure 4:**
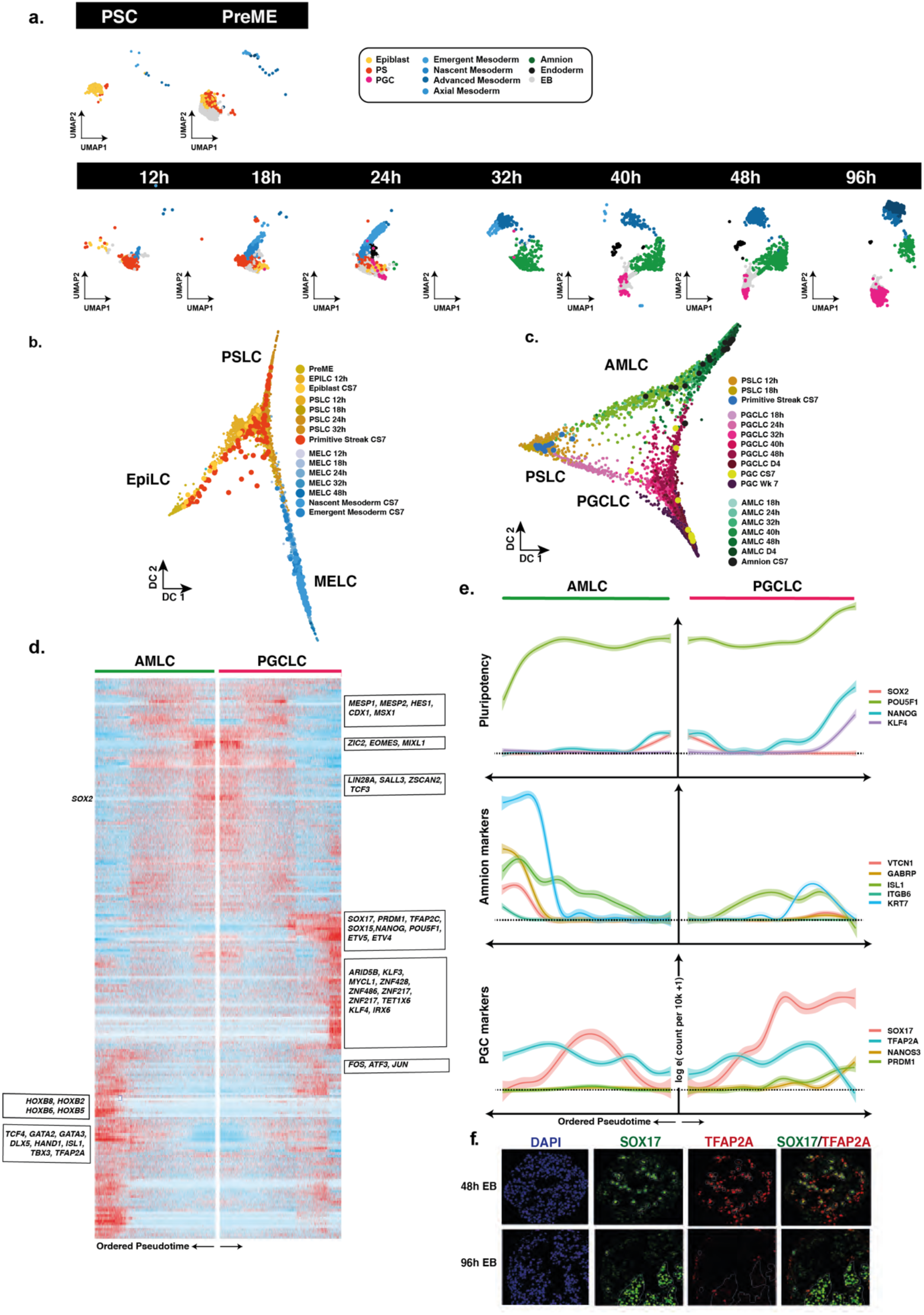
Resolving the dynamics of bifurcations in embryoid bodies. (a) Visualisation of data separated by sample time with cells annotated by transfer of labels from the human CS7 gastrula^1^; representation suggests EBs develops first through a primitive streak-like stage, early emergence of mesoderm-like cells and primordial germ cell-like cells, followed by amnion-like cells. (b) Diffusion map representation of specific clusters reveals strong bifurcation of mesoderm from the PS-like progenitors, with the remaining PS-like cells destined for other lineages. (c) Diffusion map representation of AMLC and PGCLCs shows bifurcation from common progenitor cells, with the continued association until 48h. Superimposition of cells from the CS7 gastrula labelled as PS, amnion or PGCs shows early alignment of hPGCs to PGCLCs. (d) WOT analysis to infer progenitor-descendent relationships, identifying bifurcations of individual lineages. Heatmap representants differentially expressed genes between AMLC and PGCLC ordered by pseudotime. (e) Line plot representations of essential genes ordered by pseudotime shows early up-regulation of TFAP2A in both PGCLC and AMLCs, which is sustained in AMLC. (f) IF shows TFAP2A in early PGCLCs at 48h (SOX17/TFAP2A double-positive) is lost by 96h.

We visualised the segregation of early mesoderm from precursors with primitive-streak-like identity using a diffusion map (Fig. 4b). Cells not committed to mesoderm-fate are instead predominately directed towards PGCLC or AMLC. Both UMAP and DM representations suggest that PGCLCs and AMLCs stem from highly similar progenitor cells (Fig. 4b). Interestingly, there remains some association between the PGCLC and AMLC branches until around 48h, with a number of cells falling between the two main branches. Visualisation of the PGCLC branch alongside samples from the CS7 human gastrula shows an overlap between the gastrula samples and our Wk7 in vivo PGCs and late in vitro PGCLCs (Fig. 4c). It is also worth noting that four other cells, initially labelled as PS in the human gastrula dataset, were also found to align to early PGCLCs and were reannotated accordingly. Together these observations strongly suggest that the CS7 gastrula contains samples of PGCs at different stages of specification and that our in vitro model captures the dynamics of this developmental trajectory at a much finer resolution. Cross comparison of CS7 PGCs with PGCLCs from various other in vitro models confirms a robust and conserved program of PGCLC-specification centred around the *SOX17*/*TFAP2C*/*PRDM1* network with consistent up-regulation of *TFAP2A* and other genes (Supplementary Fig. 5).

We quantified the dynamics of individual bifurcations by inferring lineage trajectories with Waddington-OT (Schiebinger *et al*., 2019), an optimal transport-based approach that allowed us to infer progenitor-progeny relationships between groups of cells statistically. By integrating these results with reduced dimensional representations of our time-course data, such as UMAP, DM, or PCA, we sought to identify the most likely earliest progenitors of PGC specification in our data. Using the ancestor-progeny relationships computed by WOT we inferred the broader lineages by first constructing a sparse network of clusters (Supplementary Fig. 6a,b) which were further grouped using a community-detection algorithm (see Methods). We assigned the inferred lineage identities to the single cells in these groups using broad marker gene expression patterns. As an initial check, we overlaid these WOT-inferred lineages onto our UMAP in Supplementary Fig. 6c, which demonstrated a good agreement with our earlier annotation-based lineage assignments with a high degree of correlation to our earlier cluster and marker-based annotations (Supplementary Fig. 6d). Using Waddington-OT inference, most terminal cell fates were effectively traced to 24h, with some cell groups traced to earlier stages.

Early mesoderm populations progressed from a PS-like state through a nascent-mesoderm-like state (nMELC) expressing *MESP1*/*2* and *T* to an emergent mesoderm-like state (eMELC), representing the highest levels of *MESP1*/*2* and downregulation of *T* (Supplementary Fig. 6e-f). Between 24 to 32h, a *PDGFRA* positive population emerged, aligned to advanced mesoderm of the human gastrula (denoted advanced mesoderm-like cells; aMELC), concomitant with the gradual loss of nMLC and eMLC subpopulations. By ordering gene expression along a diffusion pseudotime analysis, we observed the late up-regulation of several advanced mesoderm markers, *HAND1, SNAI2* and *GATA6* (Supplementary Fig. 6e-g). As the earliest specified fate, nascent and emergent mesoderm cells express several genes that may influence the balance of fates within the embryoid body, including *BMP4, WNT5A*, and *CER1*, and extra-cellular matrix genes (see e.g., Supplementary Fig. 4).

From 24h to 32h, a limited pool of *SOX17*-positive endoderm-like cells bifurcated from the PS-like subpopulation and showed sustained *NODAL* expression with subsequent upregulation of endoderm markers *FOXA1/2* (Supplementary Fig. 6h,i). Although the number of cells in this population appeared to be fewer than that of other cell lineages, it was nevertheless a conserved feature across in vitro models.

Around the 18-hour mark, the earliest PGCLCs bifurcated from a progenitor population with strong up-regulation of *SOX17, TFAP2C*, and *PRDM1* (see Supplementary Fig. 2g-h) and subsequent expression of *NANOS3*. PGCLCs also showed up-regulation of *WNT2* with early PGCLCs expressing *NODAL* (Supplementary Fig. 4).

Indeed, a comparison of PGCLC-precursor cells in high and low PGC-competence cell lines (Chen *et al*., 2019) revealed *NODAL* to be differentially expressed, consistent with a recently observed role for *NODAL* in PGCLC specification (Jo *et al*., 2021). Slightly later, at 24h, an AMLC branch also became evident, expressing *TFAP2A* and, at later time points, *ISL1*, a LIM/homeodomain transcription factor protein recently identified as an amnion marker (Guo *et al*., 2020; Yang *et al*., 2021) (Supplementary Fig. 2c-d). This AMLC branch shows an expression of *WNT6* (Supplementary Fig. 4). We identified differentially expressed genes along the separate AMLC and PGCLC lineages using the diffusion pseudotime ordering of single cells (Fig. 4d; see Supplementary Materials). Within these pseudotime trajectories, we observed that both AMLC and PGCLC showed early coordinated expression of *EOMES, MIXL1* and *ZIC*, together with rapid downregulation of *SOX2*. Moreover, we observed late expression of *VTCN1, GATA3, GATA2, ISL1* and *HAND1* in AMLCs, while the PGCLC trajectory showed late expression of PGC markers *SOX17, PRDM1, TFAP2C, SOX15, KLF4, LIN28*, and *POU5F1*. Fig. 4e shows the divergent expression patterns of crucial TFs over pseudotime to trace their rise and fall to AMLC versus PGCLC trajectories. We note an initial up-regulation of *SOX17* in AMLC and PGCLCs that is transient in AMLC but sustained in PGCLCs. Surprisingly, *TFAP2A*, which is generally considered a trophoblast or amnion marker (Krendl *et al*., 2017; Zheng *et al*., 2019; Minn *et al*., 2020), precedes *SOX17* expression and is transiently co-expressed with *SOX17* in the PGCLC trajectory. While AMLCs maintain *TFAP2A* expression, there is downregulation in PGCLCs, which was confirmed by immunofluorescence staining at the protein level (Fig. 4f). Staining of EBs for *TFAP2A* and *SOX17* confirmed their co-expression at early time points, whereas, in the 96h EB, *TFAP2A* expression is exclusive to AMLC and *SOX17* to PGCLCs and DELCs. These results, taken together, highlight the complex dynamics of PGCLC specification within our model system and identify several putative markers of specification. The most interesting was the early and transient expression of *TFAP2A* in PGCLCs. TFAP2A is an early BMP response gene that shares the TF binding site with TFAP2C (Krendl *et al*., 2017). Given that we previously found TFAP2 motifs around PGC-related genes (Tang *et al*., 2022), and that the TFAP2 family can play complementary roles, it is interesting to see if TFAP2A plays a role in PGCLC specification before the onset of TFAP2C expression.

### TFAP2A is the most upstream crucial regulator of PGC specification

To determine whether the transient *TFAP2A* expression has a role in PGC fate, we induced PGCLCs via PreME states using PSCs with a knockout mutation in *TFAP2A* and compared the outcome with the parental PSC line (Krendl *et al*., 2017) (Fig. 5a). We observed a reduction in PGCLCs in *TFAP2A* mutant cells compared to parental controls by FACS using antibodies for PGC-surface markers PDPN and AP (2.78% vs 9.28%) (Fig. 5b). Quantification of four independent experiments showed a consistent and statistically significant reduction in PGCLC specification in TFAP2A KO EBs (Fig. 5c) confirmed by immunofluorescence staining of d4 EBs generated from TFAP2A knockout cells, compared to parental control (Fig. 5d).

**Figure 5:**
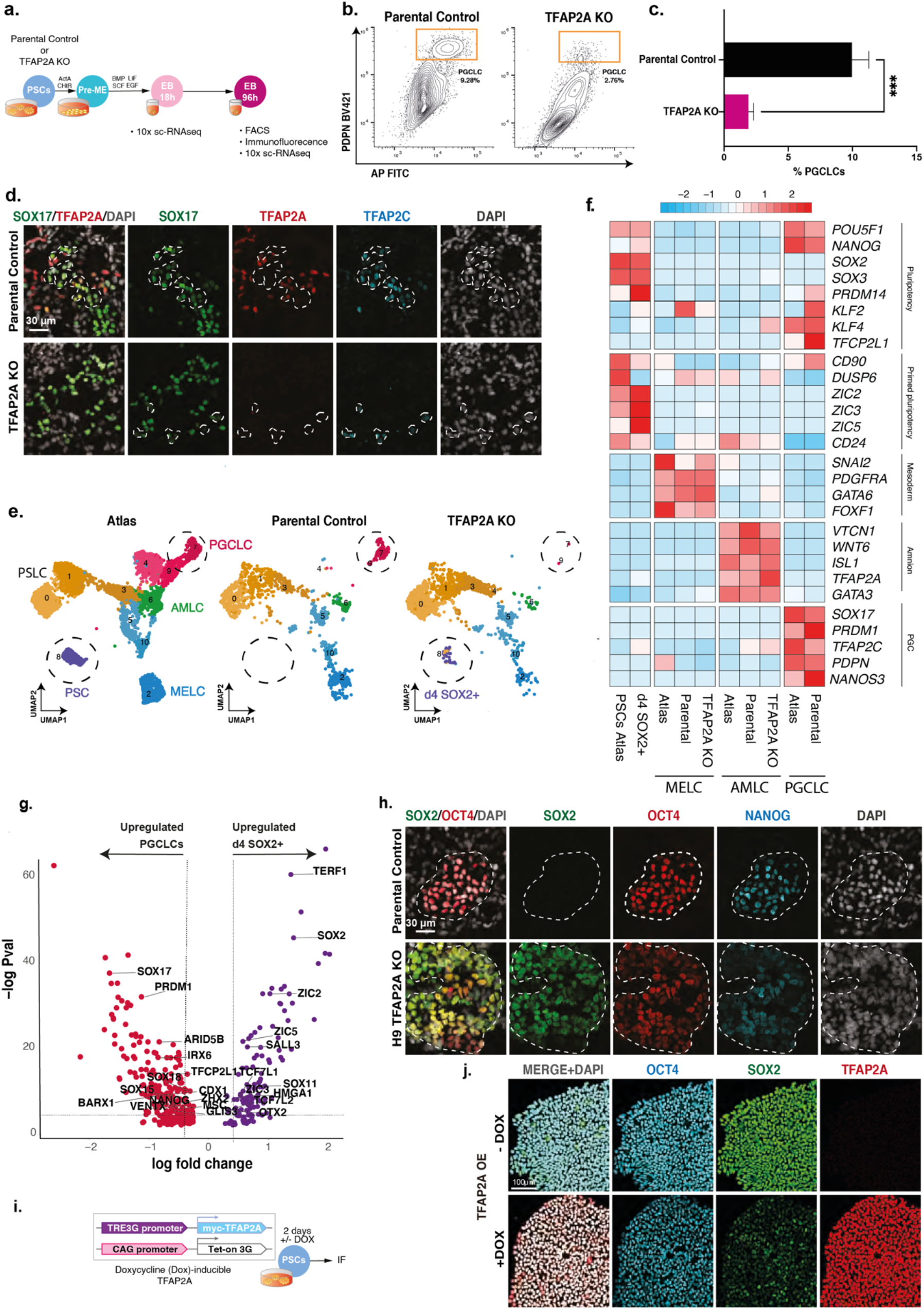
TFAP2A is a regulator of PGCLC fate. (a) Testing the role of TFAP2A in PGC specification. (b) FACS plot reveals % PGCLCs in TFAP2A KO EBs and WT parental control. (c) Immunofluorescence shows co-expression for SOX17, TFAP2A and TFAP2C in d4 EB. (d) Quantification of PGCLC (%) in WT and TFAP2A KO. (e) Aligned UMAPs for the reference data versus parental control and H9 TFAP2A KO. (f) Row-normalised gene expression demonstrate consistent expression in AMLC and MELC in the TFAP2A KO line. D4 SOX2+ cells shows expression of pluripotency genes. (g) Immunofluorescence of d4 parental EBs shows OCT4 NANOG double-positive cells (PGCLCs) but not in TFAP2A KO EBs; instead, there are OCT4, NANOG, and SOX2 triple positive cells. (h) Volcano plot for differentially expressed genes between the d4 SOX2+ cluster in TFAP2A KO vs PGCLCs in parental control) (i) Testing the role of TFAP2A overexpression in PSCs. (j) Immunofluorescence for OCT4, SOX2, TFAP2A in PSCs.

To characterise the phenotype due to TFAP2A loss of function further, we generated 10X scRNA-seq datasets for two-time points: 18h, just before the diversification of distinct lineages in embryoid bodies, and at 96h, when terminal cell fates have been established. We integrated these time points with our existing EB dataset containing all cell types for reference using Seurat. For this alignment, we generated a new clustering visualised on a UMAP in Fig. 5e.

Embryoid bodies in parental lines showed the precise formation of a MELC expressing *PDGFRA*, amnion expressing *VTCN1*, and PGCLCs expressing *NANOS3* by 96h (Fig. 5e), suggesting conserved terminal behaviour with previous lines. While there were no detectable DELCs in either the parental or TFAP2A KO line by scRNA-seq, likely due to their limited cell numbers, immunofluorescence analysis shows that rare SOX17, FOXA2 double-positive cells were present in the EB (Supplementary Fig. 7a). On the other hand, in EBs with TFAP2A KO cells, PGCLC lineages were virtually absent (Fig. 5E, Supplementary Fig. 7a-b), but aMELC and AMLC populations were present. While the TFAP2A KO appeared to lack PGCLCs, we observed a new subpopulation of cells at 96h clustered alongside pluripotent cells (Fig. 5e). This population, absent in the parental line and rare at the 18h mark in the KO line, showed expression of *SOX2* and other pluripotency markers (hereafter referred to as SOX2+d4 cells; Supplementary Fig. 7c).

To help establish the authenticity of the other fates, we generated a cross-correlation heatmap (Supplementary Fig. 7d). The SOX2+d4 cluster is most similar to PSCs in the reference population. AMLCs in the KO cluster were highly similar to AMLCs in the parental line and the reference line, with MELCs also showing consistency across all cell lines. Together these observations suggest no significant effect of *TFAP2A* loss of function for MELCs or AMLCs specification. Immunofluorescence analysis confirmed the presence of AMLCs (GATA3+ HAND1+), MELCs (HAND1+) (Supplementary Fig. 7e), and DELC cells (SOX17+, FOXA2+) in TFAP2A KO in TFAP2A KO EBs but with a minimal number of PGCLCs (SOX17+, OCT4+) (Supplementary Fig. 7a), confirming that TFAP2A had no significant effect on the other fates of the EB. We, therefore, focused on PGCLCs and the SOX2+ population.

Differential expression analysis of the SOX2+d4 population compared to parental-line PGCLCs showed that the SOX2+d4 cells expressed pluripotency and neural-plate factors, *ZIC2, ZIC5, SOX11, OTX2*, while PGCLCs showed expression of germ cell markers *SOX17, PRDM1*, S*OX15, ARID5B, TFCP2L1* and *VENTX* (Fig. 5g). We found upregulation of na*ve markers of pluripotency and neuronal lineage-associated genes in the SOX2+d4 population compared to PSCs in the reference atlas; markers included *PRDM14, KLF4, KLF6, and TFAP2C*, and neuro-related genes *ZIC2, ZNF292, FOXN3, POU3F1, SOX11, SOX4, ZIC5*, and *SALL3* (Supplementary Fig. 7f).

To validate our findings at the protein level, we performed immunofluorescence staining at d4 EBs and found expression of *SOX2* in TFAP2A KO cells even after four days of cytokine exposure. There was a rapid downregulation of *SOX2* upon BMP exposure (Supplementary Fig. 7c) in WT cells, which is critical for efficient PGCLC specification(Lin *et al*., 2014). SOX2+d4 cells also showed co-expression of OCT4 and NANOG (Fig. 5h).

We investigated if TFAP2A could potentially target SOX2 for downregulation based on these results. For this, we generated a stable dox-inducible TFAP2A PSCs line. Upon doxycycline induction of *TFAP2A* in PSCs cultured in E8 medium by dox for two days (Fig. 5i), we observed a substantial reduction in SOX2 levels after TFAP2A overexpression by immunofluorescence (Fig. 5j). *POU5F1* was also slightly reduced. Together, our results suggest that TFAP2A is a regulator of PGCLC fate and may participate in the downregulation of SOX2 and other targets impeding PGCLC specification.

## Discussion

In vitro models have been of vital importance for unravelling the transcriptional network responsible for human germ cell competence and specification (Teo *et al*., 2011; Irie *et al*., 2015; Sasaki *et al*., 2015; Chen *et al*., 2017; Kobayashi *et al*., 2017; Kojima *et al*., 2017, 2021; Pierson Smela *et al*., 2019; Sybirna *et al*., 2020). In this study, we characterise in vitro models for the derivation of PGCLCs from PSCs by highly resolved single-cell transcriptomics and comprehensive comparison to in vivo references in human, non-human primates, and other in vitro models of gastrulation.

Notably, we found that PGC competent PreME cells exist transiently within a continuum of states extending from PSCs to mesendoderm (ME). Our analysis showed that clusters enriched for PGC-competent populations present a particular signalling signature, characterised by active Nodal and WNT signalling. There is low expression of BMP inhibitors (*BAMBI* and *CER1*) in competent cells compared to the ME-dominated cluster with the highest levels in non-competent cluster 22, which likely impedes PGC specification. *BAMBI* is a direct target of WNT signalling (Sekiya *et al*., 2004), while activation of *CER1* occurs via both WNT and Nodal signalling (Katoh and Katoh, 2006; Martyn, Brivanlou and Siggia, 2019). PGCLC-competent clusters also show transient downregulation of *OTX2* and higher levels of *NANOG* compared to non-competent clusters, which we recently found is conducive to transition to the PGCLC state (Tang *et al*., 2022). Concomitantly, there is an increase in the levels of *EOMES*, which has a prominent role in human PGC-competence (Chen *et al*., 2017; Kojima *et al*., 2017), but further activation of mesoderm factors hinders PGC specification. The tight signalling axis, transcription factor levels and intrinsic heterogeneity modulating competence are consistent with a relatively small number (∼100-200) of founder PGCs in vivo (Saitou, Barton and Surani, 2002; Kobayashi *et al*., 2017).

Specification of PGCLCs in vitro occurs within a 3D aggregate that consists of a hitherto poorly characterized fraction of somatic components. Currently, PGCLCs can be induced in various 2D aggregates but more efficiently in 3D embryoids, highlighting the importance of the structure, cell-cell interactions or signalling from adjacent tissues (Minn *et al*., 2020, 2021). Here we have shown that these somatic cells collectively represent those in the posterior region of the embryo during gastrulation. Among the somatic cell types, we note the early formation of mesoderm-like cells, which display strong expression of BMP, WNT, and ECM components that may be important for PGC-fate and potentially play a similar role to that of extraembryonic mesoderm in the embryo, and endoderm-like cells that are double positive for *FOXA2/SOX17*. Furthermore, we also observe the emergence of *ISL1*/*VTCN1* expressing amnion cells, providing evidence that amnion formation continues from the posterior epiblast during gastrulation, as recently suggested in a study on marmoset (Bergmann *et al*., 2022).

Mapping the cells to a 3D primate embryo showed that PSCs best correspond to the anterior region of the embryonic disc, while PreME cells shifted towards the posterior end. Conversely, cells within the newly formed embryoid body at 12h, which transcriptionally resemble a primitive streak, mapped best to the posterior end of the embryonic disc, with PGCLCs mapping to a *SOX17*/*TFAP2C*/*NANOS3* positive region at the boundary between the posterior-most epiblast and amnion.

The origin of human PGCs remains unresolved due to the inaccessibility of human embryos, but bilaminar disc embryos from other species provide valuable information. In species such as the rabbit and pig, PGCs originate from the posterior epiblast, but the amnion develops later, indicating that the development of the amnion and PGCs in some cases are temporally unconnected. In humans and non-human primates, development of the amnion commences prior to PGC specification, but according to our work and by others (Bergmann *et al*., 2022; Rostovskaya *et al*., 2022) amniotic cells continue to emerge later from the posterior epiblast, co-incidentally with the specification of PGCs at the time of primitive streak formation. In cynomolgus monkeys, the earliest PGCs have been reported in the amnion, with the majority found later in the epiblast. One possibility is that these early PGCs may arise from intermediate cells that are en route to the amnion but are but not fully committed as squamous amniotic epithelium as observed in our data (Fig. 4c). To contribute to the founder PGC pool, PGCs arising in the amnion would need to migrate against the continuing amnion growth. We posit that at this stage of development in humans and non-human primates, amnion cells continue to be specified with nascent PGCs arising at the posterior-most end of the epiblast during the early PS stage.

In our model, AMLC and PGCLC progenitors display early expression of *TFAP2A*, a pioneer factor previously associated with the amnion (Shao, Taniguchi, Townshend, *et al*., 2017). Whilst there is subsequent downregulation of *TFAP2A* in PGCLCs, expression is sustained in the amnion. Surprisingly, the knockout of *TFAP2A* did not have a detectable effect on AMLCs, which merits further investigation, but notably resulted in an almost complete abrogation of PGCLCs.

In PGCLCs, TFAP2A is rapidly replaced by the expression of TFAP2C, suggesting otherwise mutually exclusive expression after a brief window of co-expression. Interestingly, TFAP2A shares the same transcription factor binding motif as TFAP2C (Krendl *et al*., 2017). TFAP2C is essential for PGC development (Kojima *et al*., 2017) and acts as both an activator and a repressor during PGC specification but it is not sufficient for PGC fate in the absence of cytokines (Kobayashi *et al*., 2017). In the PGCLC pseudotime trajectories, we saw early upregulation of TFAP2A (12h), followed by expression of SOX17 and TFAP2C (18h), and later, activation of PRDM1 (24-32h) by SOX17(Tang *et al*., 2022) (Supplementary Fig. 8). In some instances, TFAP2A functions similarly to TFAP2C (Hoffman *et al*., 2007; Li and Cornell, 2007). Our work suggests that TFAP2A expression is transient but essential for initiating the PGC transcriptional network, and may directly or indirectly repress *SOX2* and other factors.

TFAP2A KO EBs show an emergent population (SOX2+ d4 cells) found to align to pluripotent stem cells, with the expression of the core pluripotency genes; *SOX2, POU5F1* and *NANOG*. Differential gene expression between PSCs and SOX2+ d4 cells shows aberrant upregulation of na*ve markers *KLF4, TFAP2C* and *PRDM14* and genes associated with the neuronal lineage, including *ZNF292, FOXN3, SALL3, ZIC2, POU3F1* in SOX2+ d4 cells.

There is rapid downregulation of SOX2 during human PGCLC-induction (Kobayashi *et al*., 2017); indeed, sustained SOX2 expression prevents PGCLC specification due to elevated differentiation into the neuronal lineage (Lin *et al*., 2014), which could in part explain the expression of related neuronal markers in the TFAP2A mutant cells. The combinatorial role of SOX17-OCT4 involved in human germ cell fate (Tang *et al*., 2022) might benefit from a repression of SOX2 to favour the SOX17-OCT4 interaction on the compressed motif.

We provide insight into early human development with the transient emergence of the germ cell competent PreME cells in a model mimicking human gastrulation starting with PSC. Our study suggests continuing emergence of the amnion from the posterior epiblast at the time of PGC specification during early gastrulation; the amnion and PGC likely arise from highly similar progenitor exemplified by TFAP2A expression. The loss of function has a marked effect on PGC specification but without a detectable effect on the amnion. Accordingly, PGCs likely emerge from the posterior epiblast predominantly, notwithstanding a sub-set in the early amnion (Fig. 6). Of great interest would be to test, when possible, the predictions we make by direct observations in extended cultures of developing human embryos.

**Figure 6:**
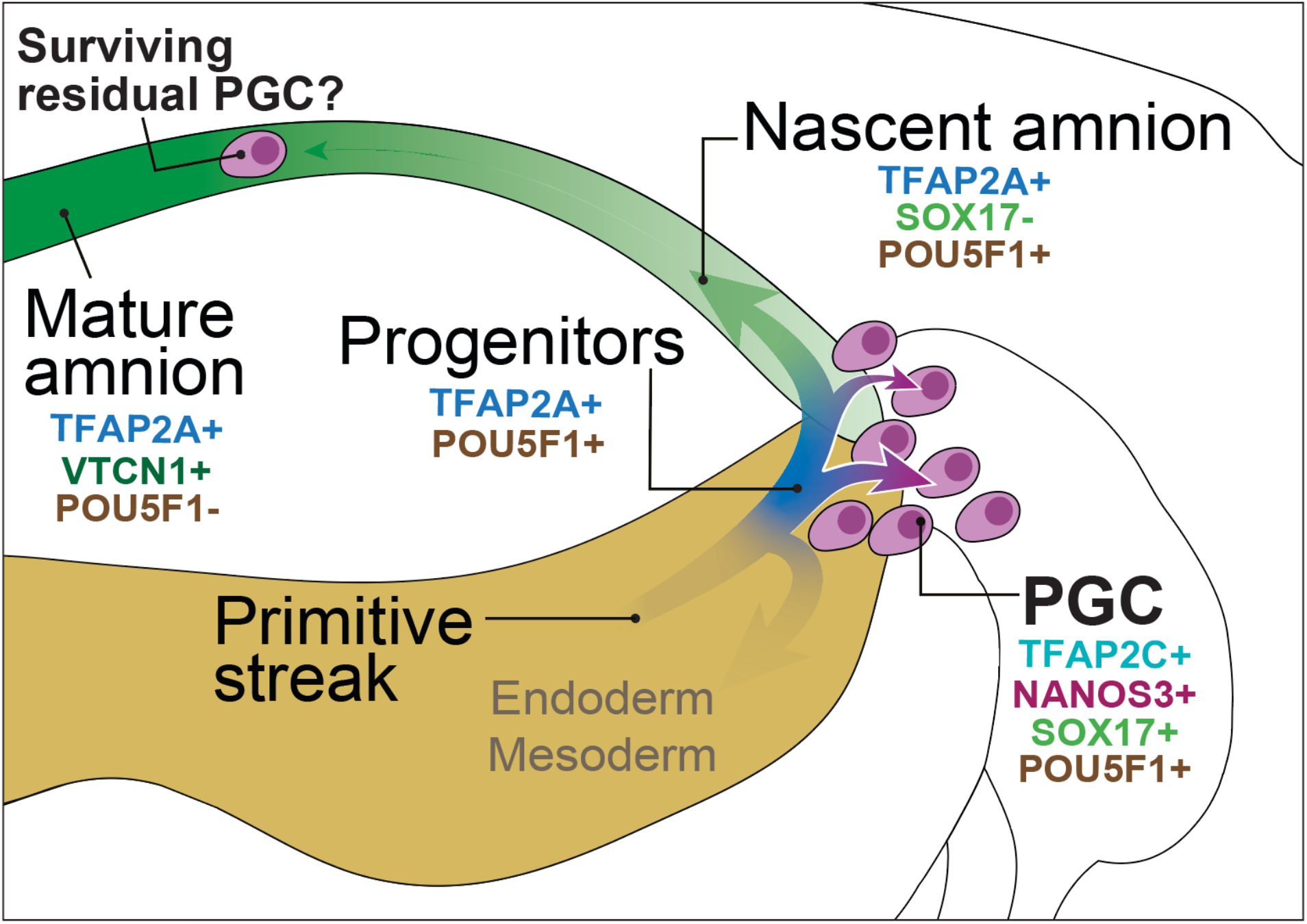
A unifying model of PGC specification in bilaminar disc embryos. PGCs are specified from *TFAP2A*-positive progenitors at the posterior end of the embryonic disc, which also give rise to nascent amnion. PGCs in the amnion specified at an earlier stage might only contribute to the founder PGC pool if they can migrate against the flow of nascent amnion expansion.

## Material and methods

### Cell culture

H1 NANOS3-tdTomato PSC line was previously generated in the lab (Kobayashi et al., 2017). H9 parental and TFAP2A KO cells were kindly provided by Micha Drukker (Krendl et al., 2017). All cell lines were confirmed as mycoplasma negative. PSCs were maintained on vitronectin-coated plates in Essential 8 medium (Thermo Fisher Scientific) according to the manufacturer’s protocol. Cells were passaged every three to four days using 0.5 mM EDTA in PBS without breaking cell clumps.

For the 4i condition, undifferentiated PSC cells were maintained on irradiated mouse embryonic fibroblasts (MEFs) (GlobalStem) in 4i medium (Irie et al., 2015). 4i were passaged every three to five days using TrypLE Express (Gibco) quenching with MEF media and filtered with 50 μm cell filter (PERTEC). ROCK inhibitor (10 μM; Y-27632, TOCRIS Bioscience) was kept in the culture for 24 h after passaging.

Mesendoderm induction was performed as reported in (Kobayashi et al., 2017). PSCs were dissociated into single cells using TrypLE Express and seeded onto vitronectin coated plates at 500,000 cells per well of 6-well plate and cultured in mesendoderm (ME) induction medium for 10 to 12 hours. ME medium is based on aRB27

Primordial germ cells were induced as reported previously (Irie et al., 2015; Kobayashi et al., 2017). For this PreME cells were disaggregated into single cell solution using TrypLE, then 4,000 cells per well were seeded into ultra-low attachment 96-well plates (Corning Costar) in PGC induction medium. Mesendoderm, PGCLC and definitive endoderm were induced from NANOS3–tdTomato reporter PSCs as described before (Kobayashi et al., 2017) using the aRB27 basal medium, which was composed of Advanced RPMI 1640 Medium (Thermo Fisher Scientific) supplemented with 1% B27 supplement (Thermo Fisher Scientific), 0.1 mM NEAA, 100 U/ml penicillin, 0.1 mg/ml streptomycin, 2 mM L-glutamine. To induce mesendoderm, trypsinized hPSCs were seeded on vitronectin-coated dishes at 200,000 cells per well in a 12-well plates and cultured in mesendoderm induction medium for 12 (PreME) and 24 (ME) hours. Mesendoderm induction medium contained aRB27 medium supplemented with 100 ng/ml activin A (Department of Biochemistry, University of Cambridge), 3 μM GSK3i (Miltenyi Biotec) and 10 μM of ROCKi (Y-27632, Tocris bioscience). To induce definitive endoderm from ME, mesendoderm induction medium was replaced with definitive endoderm induction medium after washing with PBS once and cells were cultured for a further 2 days. Definitive endoderm induction medium was composed of aRB27 medium supplemented with 100 ng/ ml activin A (Department of Biochemistry) and 0.5 μM BMPi (LDN193189, Sigma).

To induce PGCLCs, PreME cells were trypsinized into single cells and harvested into Corning Costar Ultra-Low attachment multiwell 96-well plate (Sigma) at 4,000 cells per well in hPGCLC induction medium, which composed of aRB27 medium supplemented with 500 ng/ml BMP4,10 ng/ml human LIF (Department of Biochemistry), 100 ng/ml SCF (R&D systems), 50 ng/ml EGF (R&D Systems), 10 μM ROCKi, and 0.25% (v/v) poly-vinyl alcohol (Sigma). Cells were cultured as floating aggregate for 2-4 days.

To collect PSCs, PreME, ME, DE, PGCLCs, cells were trypsinized with 0.25% trypsin/EDTA at 37 °C for 5-15 min. DE was stained with PerCP-Cy5.5 conjugated anti-CXCR4 antibody (Biolegend). Cell suspension was subjected to FACS by SH800Z Cell Sorter (Sony) and analyzed by FlowJo software.

### Collection of human PGCs from human embryos

Human embryonic tissues were used under permission from NHS Research Ethical Committee, UK (REC Number: 96/085). Human embryonic samples were collected following medical or surgical termination of pregnancy carried out at Addenbrooke’s Hospital, Cambridge, UK with full consent from patients. Crown-rump length, anatomical features, including limb and digit development, was used to determine developmental stage of human embryos with reference to Carnegie staging (CS). The sex of embryos were determined by sex determination PCR as previously described (Bryja and Konečný, 2003).

Human embryonic genital ridges from individual embryos (wk7) were dissected in PBS and separated from surrounding mesonephric tissues. The embryonic tissues were dissociated with Collagenase IV (2.6 mg/ml) (Sigma, C5138) and DNase I (10 U/ml) in DMEM-F/12 (Gibco) at 37°C for 15-30 minutes (depending on tissue size). Tissues were pipette up and down for five times every 10 minutes to facilitate dissociation into single cell suspension. After that, samples were diluted with 1 ml FACS medium (PBS with 3% fetal calf serum & 5 mM EDTA) and centrifuged at 500 xg for 5 minutes. Cell pellet was suspended with FACS medium and incubated with 5 μl of Alexa Fluor 488-conjugated anti-alkaline phosphatase (BD Pharmingen, 561495) and 5 μl of APC-conjugated anti-c-KIT (Invitrogen, CD11705) antibodies for 20 minutes at room temperature with rotation at 10 revolutions per minutes (rpm) in dark. Cell suspension was then diluted in 1 ml FACS medium and centrifuged at 500 xg for 5 minutes. After removing the supernatant, the cell pellet was resuspended in FACS medium and passed through a 35 µm cell strainer. FACS was performed with SH800Z Cell Sorter (Sony) and FACS plots were generated by FlowJo software.

### Fluorescence-activated cell sorting (FACS)

PSCs, 4i, PreME and ME cells were harvested using TrypLE (GIBCO) at 37°C for 2-3 min. Embryoid bodies were collected and dissociated into single cells using Trypsin-EDTA solution 0.25% at 37°C for 5 to 15 min. Dissociated cells were washed and resuspended in the FACS buffer (PBS 3% FCS). DE samples were stained with PerCP-Cy5.5 conjugated anti-CXCR4 antibody (Biolegend) for 1h on ice. Samples were washed with PBS, stained with DAPI (1:10,000) and sorted on a SONY SH800 sorter.

Human embryonic genital ridge or mesonephros from a week 7.0 male embryo were collected in dissection medium (DMEM (Gibco), 10% FCS, 1 mM sodium pyruvate (Sigma)). Embryonic tissues were dissociated with 300 μL collagenase IV (2.6 mg/mL in DMEM-F/12) supplemented with DNaseI (10U/mL) per genital ridge and incubated for 10 minutes at 37°C with mixing by pipetting up and down. Then, cells were washed with 1 mL FACS buffer (PBS with 3% FCS and 5 mM EDTA). Resuspended with 75 μL FACS buffer and stained with 0.5 μL alexa Fluor 488-conjugated anti-alkaline phosphatase (BD Pharmingen, 561495) and 25 μL of PerCP-Cy5.5-conjugated anti-CD117 (BD Pharmingen 333950) for 15 minutes at room temperature. Samples were washed with PBS and sorted on a SONY SH800 cytometer. Flow cytometry data was analysed on FlowJo v10 (FlowJo LLC).

### Immunofluorescence

Embryoid bodies (EBs) were fixed in 4% PFA for 2h at 4 °C and embedded in O.C.T. compound (Cellpath) for frozen sections. Each sample was incubated with primary antibodies for 1–2 h at room temperature or overnight at 4°C and then with fluorescent-conjugated secondary antibodies and DAPI (Sigma) for 1 h at room temperature. Samples were then imaged under a Leica SP8 upright or inverted scanning confocal microscope.

Cells were cultured on ibidi µ-Slide and fixed in 4% PFA for 30 minutes at 4°C. Embryoid bodies were fixed in 4% PFA for 2 hours at 4°C and embedded in OCT compound for frozen sections. The samples were incubated with primary antibodies overnight at 4°C and subsequently with fluorescence-conjugated secondary antibodies (Thermo Fisher Scientific) and DAPI for 1 hour at RT. The primary antibodies used are: anti-GFP (abcam, ab13970), anti-PRDM1 (Cell Signaling Technology, 9115), anti-SOX17 (R&D, AF1924), anti-TFAP2C (Santa Cruz Biotechnology, sc-8977), and anti-OCT4 (BD Biosciences, 611203). Samples were imaged under Leica SP8 upright or inverted scanning confocal microscope.

### 10X genomics

For each stage, 5,000 cells were sorted into an eppendorf tube containing PBS with 0.04% weight/volume BSA (400 μg/mL). Samples collected are listed in table 2.6. During sorting, dead cells, debris and doublets were gated out. Sorted cells were directly taken for 10x processing at Cancer Research UK, Cambridge Institute and loaded into the 10x-Genomics Chromium using the single cell 3’ reagents kit v2. Libraries were prepared as per the manufacturer’s instructions and pooled for sequencing so that all lines would include all samples. Libraries were sequenced, aiming at a minimum coverage of 50,000 raw reads per cell, on an Illumina HiSeq 4000 (paired-end; read 1: 26 cycles; i7 index: 8 cycles, i5 index: 0 cycles; read 2: 98 cycles).

### Bioinformatics

#### 10X RNA sequencing processing

Multiplexed single-cell libraries were processed using the 10X Genomics cell ranger pipeline. Reads were aligned to a reference genome (Homo sapiens GrCh38) using STAR (Dobin et al., 2013), and quantification of genes against an annotation reference (based on Ensembl GrCh38 v90).

#### Analysis

Initial analysis of our data was done using Seurat (v3.1.4) (Stuart et al., 2019). Count data was normalised and scaled using NormalizeData based on log counts per 10000 (logCP10k) and scaled using ScaleData. Clusters were generated using FindCluster with resolution of 0.1. Nearest neighbour graphs and UMAP plots were calculated using the first 20 PCs.

Heatmaps of gene expression were generated based on row-scaled values using pheatmap (v.1.0.12) with cross-correlations calculated based on Pearson’s correlation and visualised using pheatmap.

#### Integrative analysis

Individual datasets were first curated to remove pre-implantation and extraembryonic tissues. Datasets were then integrated based on logCP10k using FindIntegrationMarkers with 5000 integration features and k.filter=50. Data was integrated based on CCA with 5000 features and using the first 20 PCs. Joint clustering was generated based on the integration-corrected gene expression matrices using the FindClusters function with complexity parameter uniformly incremented form 0.1-0.9 in steps of 0.1. For visualisation purposes we used parameter of 0.9 for the figures within the paper.

For initially establishing cell fates expression of key marker genes were plotted as a heatmap using pheatmap. To establish veracity of cell types between datasets, a scatter plot of differential expression was used with the x-axis showing logFC of a specific cluster vs a reference cell type/cluster (e.g., cluster 0 vs PSCs) with the y-axis showing the same comparison (cluster 0 vs PSCs) in the second dataset. Genes in the top right and bottom left quadrants represented conserved changes between the two datasets, whilst genes to the top left or bottom right represented dataset specific changes.

As a preliminary visualisation of individual bifurcations, diffusion maps were generated for selected sets of subclusters using destiny (v2.12.0) (Angerer et al., 2016) based on integration-corrected expression matrices.

#### Differential expression analysis

Unless otherwise indicated, differential expression between two groups was done in Seurat using MAST (Finak et al., 2015). For volcano plots, genes were filtered to show genes with adjusted p-values <0.05 with a >1.2 FC.

#### Mapping of cells from CS7 gastrula to embryoid bodies

Carnegie stage 7 human gastrula annotations were projected onto our EB dataset based on statistically enriched proximity in nearest neighbour graphs. Specifically, the aligned datasets were subsetted on the human CS7 gastrula and EB dataset and used to calculate a KNN graph (using the FindNeighbours function). For each cell within our EB dataset, the enrichment of individual CS7 gastrula annotations was calculated using a hypergeometric test, and final annotations assigned based on adjusted p-values. Cells that showed no significant overlap in KNN graphs were not assigned a lineage.

#### Mapping of cells to the CS6 marmoset embryo

Cells within our EB were mapped to the marmoset embryo based on proximity in KNN-graphs in the CCA aligned datasets. Aligned datasets were first subsetted on the marmoset dataset and EB dataset. For a cell, *j*, in the EB dataset, we calculated the KNN from the CS6 embryo, with positions at positions {***r***_**1**_, ***r***_**2**_,.., ***r***_*k*_: ***r***_*i*_ ∈ ℝ^3^}, and calculated the shared nearest neighbour (SNN) vector ***θ***^(*j*)^ = {*θ*_**1**_, *θ*_**2**_,.., *θ*_*k*_}. Weights were normalised 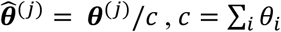 and a projection of cell *j* calculated as: 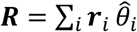, where ***r***_*j*_ ∈ ℝ^3^ denotes a 3-dimensional position vector of marmoset cell *j*. After mapping of individual cells, the density of specific groups e.g., PGCLCs (cluster 5 and 6), AMELC (clusters 7, 9 and 10), basal (cluster 1), was calculated using the MATLAB function mvksdensity.

#### Doublet detection

To minimise doublets in our analyses, we limited the number of cells loaded into each chip, with each sample capturing around 1000-2000 cells. Potential doublets were identified computationally for each individual sample using the R package DoubletFinder(McGinnis, Murrow and Gartner, 2019). For samples with ∼1000 captured cells we assumed a doublet rate of 1%, and for samples with ∼2000 cells we assumed a 2% doublet rate. No cluster analysed in this paper was found to contain a high level of doublets.

#### Waddington Optimal Transport analysis

Highly variable genes were computed across all single PSCs, PreME and EB cells, and used as input to PCA, with the first 50 PCs computed using irlba. Cells were assigned to clusters as described above, which were used as the basis for WOT. Transport maps were computed with parameters (λ1=1, λ2=50, ε=0.01) between all pairs of time points using the PSCs as 0hours, PreME as 12 hours, and all subsequent time points as 12+ti for *i* ∈ {1, 2, …, *T*} and *T* = {12, 18, 24, 32, 40, 48, 96}. Ancestor contributions to populations at subsequent time points were estimated from these transport maps using the OT trajectory command-line interface (CLI) function. Cell mass contributions between clusters across time points were concatenated into a cluster:timepoint X cluster:timepoint matrix, where the rows denote the contribution of clusterj timepointi to clusterj timepointi+1. A power threshold (p=30) was used to enforce sparsity on this matrix with values ≤ 0.1 censored to 0. This sparse matrix was then used as a weighted adjacency matrix to compute a directed KNN graph (k=5), as shown in Supplementary Figure 6. Meta-clusters were defined on this graph using the Walktrap community detection algorithm implemented in igraph, which were annotated based on the mean expression level of single-cells that contribute to each original cluster (Supplementary Figure 6). These annotations were then mapped back onto the original constituent single-cells based on their cluster identity.

## Acknowledgments

M.A.S. was supported by a Wellcome Investigator Awards in Science (209475/Z/17/Z), a MRC – Research Grant (RG85305) and a BBSRC – Research Grant (G103986). A.C.V. was supported by the Wellcome 4-Year PhD Programme in Stem Cell Biology and Medicine (203831/Z/16/Z) and the Cambridge Commonwealth European and International Trust. W.W.C.T. received a Croucher Postdoctoral Research Fellowship and was supported by the Isaac Newton Trust. T.K. and M.A.S. was supported by Butterfield Awards of Great Britain Sasakawa Foundation. T.K. was supported by the Astellas Foundation for Research on Metabolic Disorders. T.E.B. was supported by a Wellcome Sir Henry Dale Fellowship. Marmoset embryo research is generously supported by the Wellcome Trust (WT RG89228, WT RG9242), the Centre for Trophoblast Research, the Isaac Newton Trust and JSPS KAKENHI 15H02360, 19H05759.

We would like to thank Roger Barker and Xiaoling He for providing human embryonic tissues, and Charles Bradshaw for bioinformatic support. We also thank The Weizmann Institute of Science for the WIS2 hESC line and the Genomics Core Facility of CRUK Cambridge Institute for sequencing services. We thank members of the Surani lab for insightful comments and critical reading of the manuscript. J.C.M. acknowledges core support from EMBL and from Cancer Research UK (C9545/A29580), which supports M.D.M.

## Availability of materials

Any enquiries on reagents and cell lines can be directed to. Plasmids generated in this study will be made freely available upon request. Modified human embryonic stem cell lines generated in this study will be made available on request upon completion of a Materials Transfer Agreement.

Single cell RNA-seq (10X) data has been deposited at ArrayExpress under accession numbers E-MTAB-11283 and E-MTAB-11305. Code for repeating analyses will be available via a GitHub repository https://github.com/cap76/PGCLC.

## Author contributions

ACV, CAP, MAS wrote the manuscript with input from all authors. ACV, CAP, MDM designed experiments and performed analysis. ACV generated human data. SB and ES generated marmoset data. ACV, WWCT, TK, FCKW performed experiments. MAS, JCM, and TEB supervised.

