## Supplementary material for "Origin and segregation of the human germline": Suplementary figures and Supplemental Table 1

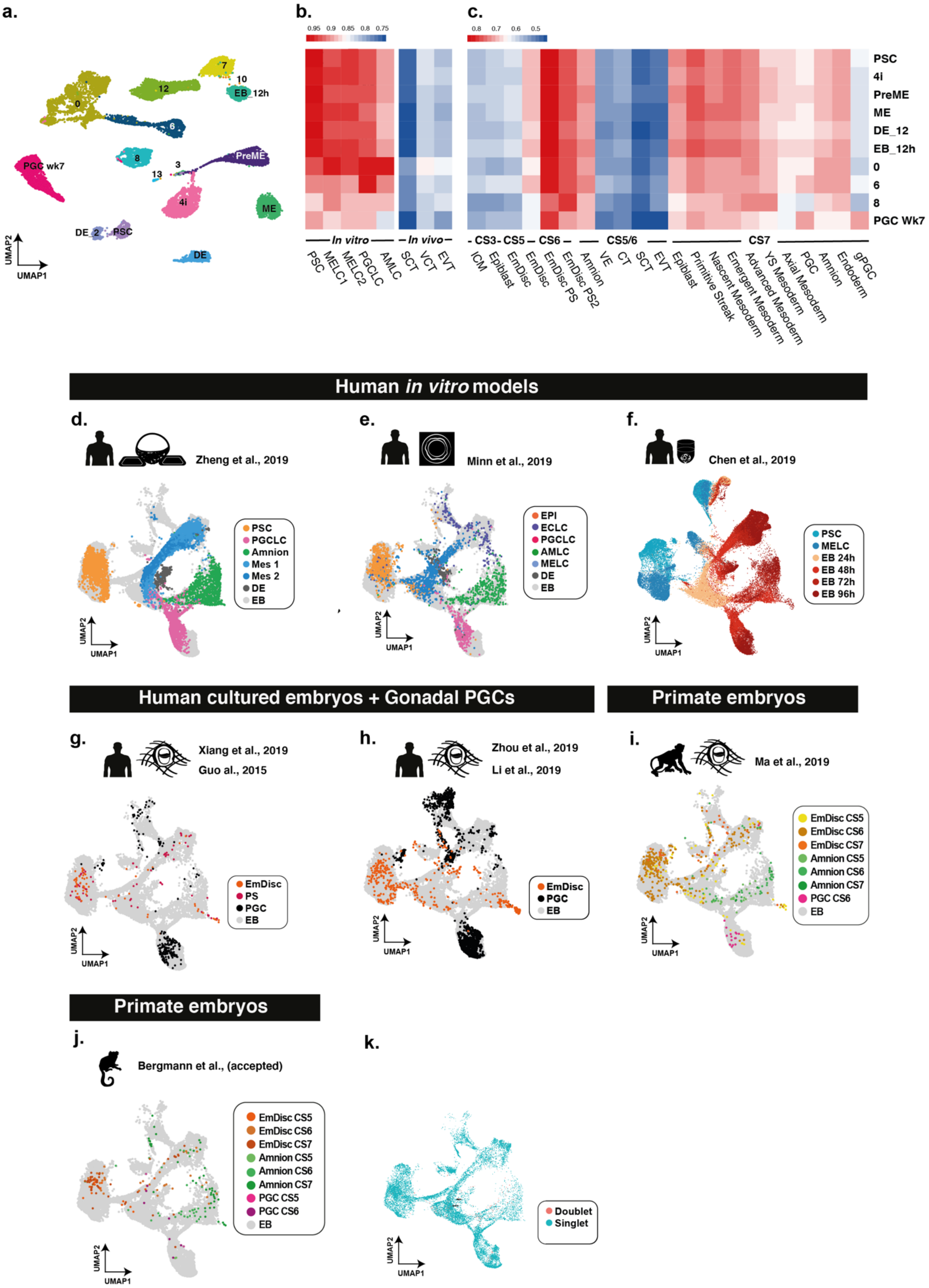


**Supplementary Figure 1**: (a) Dataset following UMAP dimensionality reduction. Louvain clustering identified 15 populations. (b-c) Pseudobulk comparison of individual clusters with existing datasets shows a high degree of correlation with embryonic lineages and amnion but less so, with extraembryonic tissues and preimplantation samples. Preliminary correlation suggests an integrative alignment with several gastrulation and germ cell datasets used to create a PGC and gastrulation roadmap, with (d-j) showing aligned samples from the individual datasets. (k) Estimation of doublet cells; pink dots – singlets blue dots.

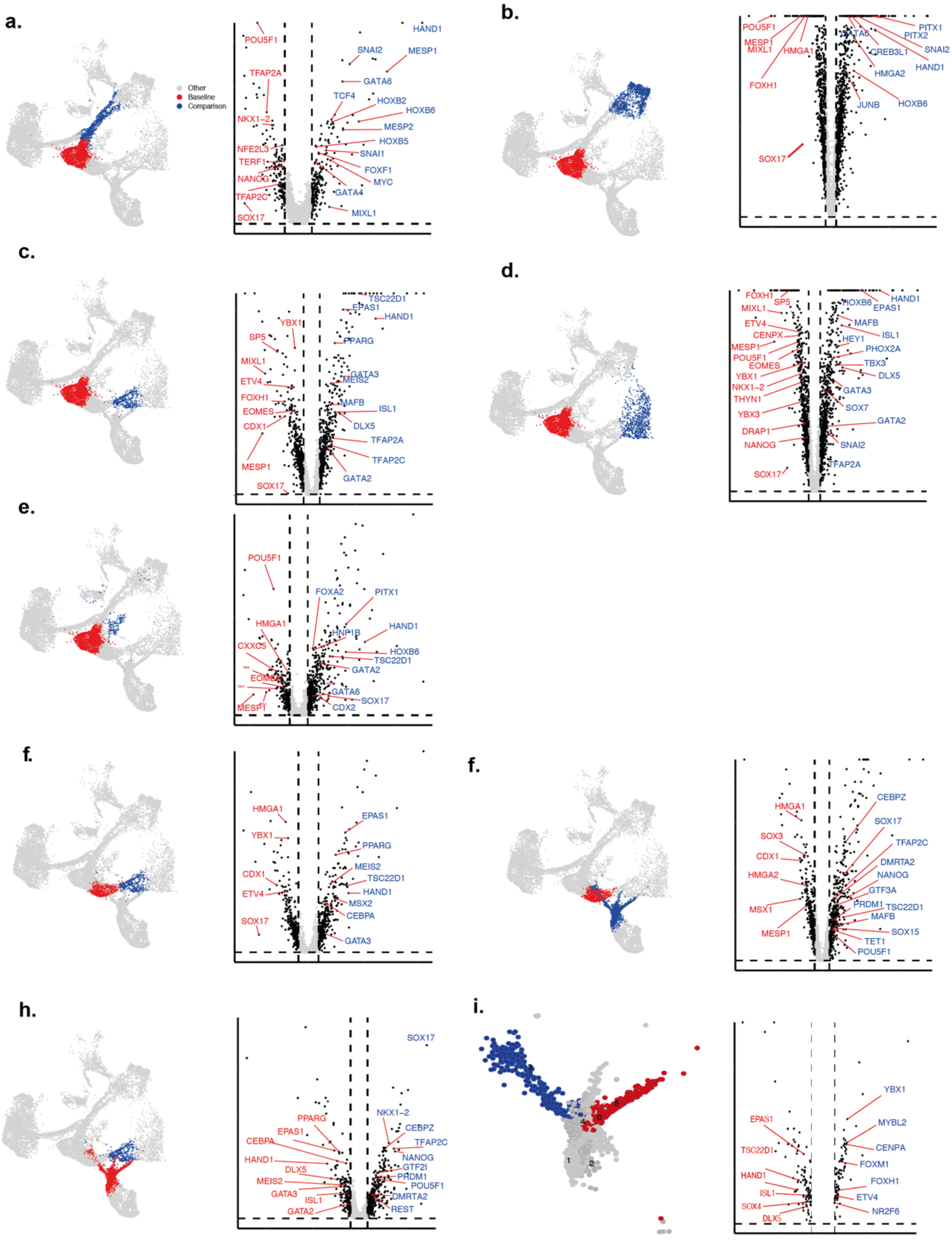


**Supplementary Figure 2**: Volcano plots showing differential expression between early (a) and late mesoderm (b), early (c) and late amnion (d), and definitive endoderm (e). Volcano plots compare progenitor for early and late amnion or PGCLC populations (f-i).

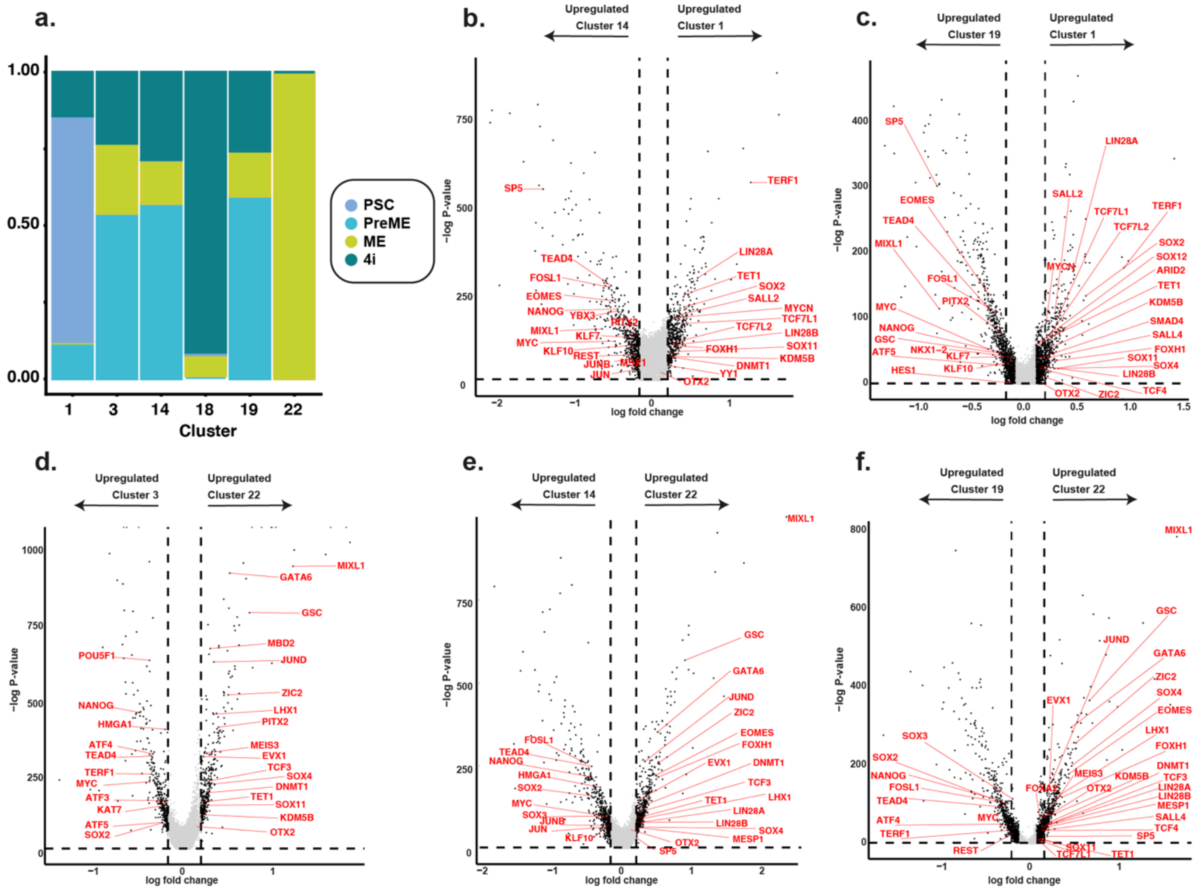


**Supplementary Figure 3**: (a) A bar plot depicting the fraction of cells from individual conditions that fall into each subcluster revealing a PSC-dominant (Cl1), ME-dominant (Cl22), and competent-dominant groups (Cl3, Cl14 Cl19). (b-f) Comparison of expression in the competent-dominant groups (cluster3, 14 and 19) with PSCs in subcluster 1 or ME cells in cluster 22 identify several putative regulators of competence.

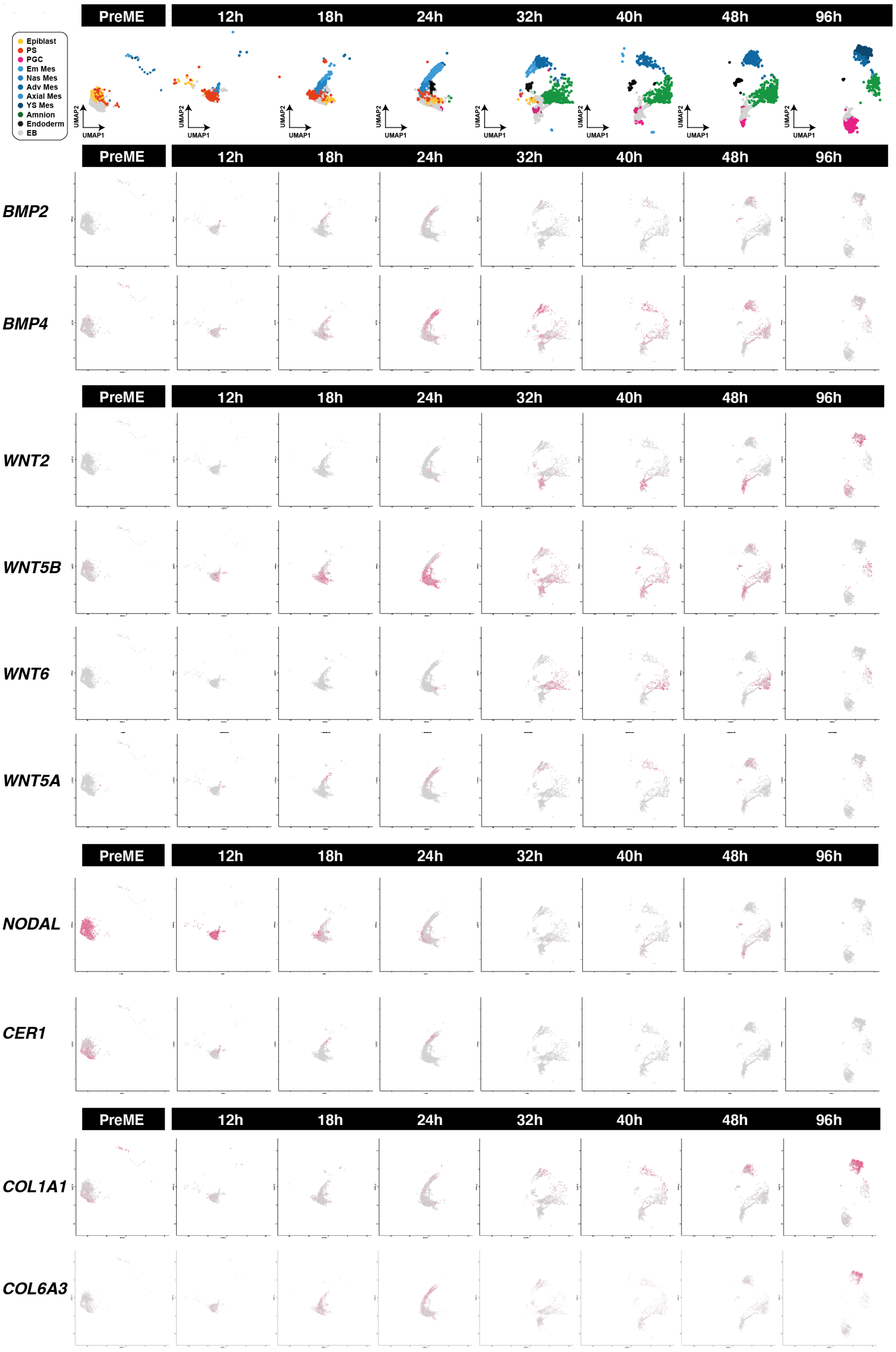


**Supplementary Figure 4**: Expression of signalling pathways show early loss of *NODAL* with later *BMP*/*WNT* signalling arising from mesoderm and amnion. Mesoderm shows expression of extracellular matrix molecules.

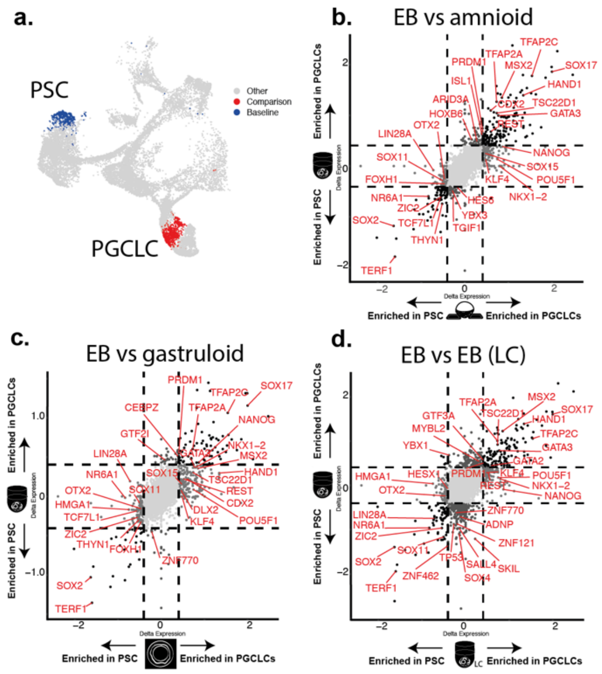


**Supplementary Figure 5**: Cross-comparison of differentially expressed genes between PGCLCs and PSCs reveals conserved markers across datasets, and highlight the robust, conserved *SOX17*/*PRDM1*/*TFAP2A*-centric network in embryoid bodies, microfluidic amnioids, and micropatterned gastruloids.

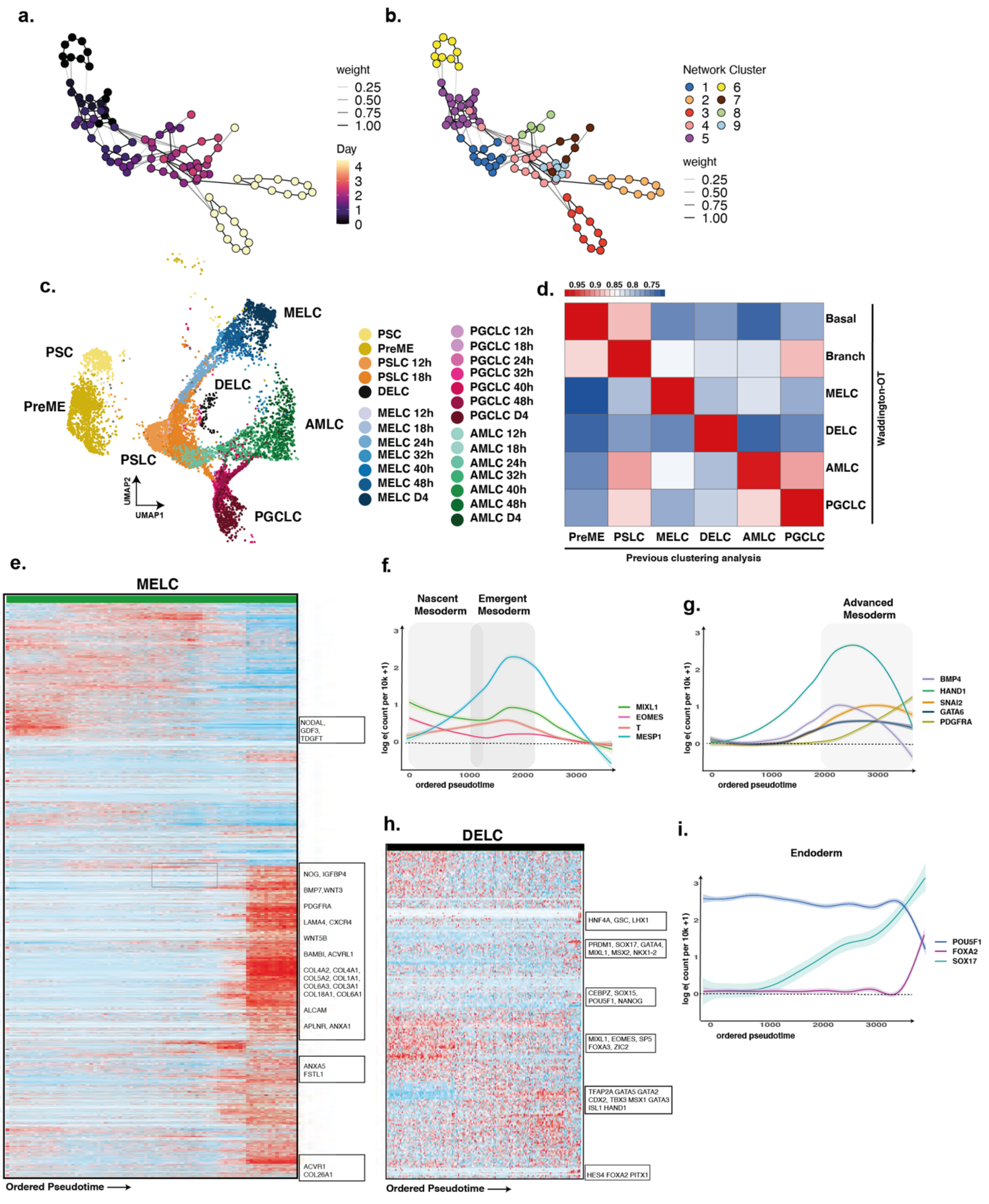


**Supplementary Figure 6**: (a-b) Waddington-OT was performed to infer lineages within the EBs. (c) Inferred WOT lineages are shown projected onto aligned UMAP representation. (d) The correlation of gene expression between WOT-inferred lineages and cluster-based lineages shows a high degree of correspondence. (e) Heatmap of gene expression in mesodermal lineages arranged by pseudotime. (f, g) Line plot representation of key genes arranged by pseudotime shows progression of mesoderm markers over time, with early up-regulation of *EOMES*/*T*/*MESP1*, and later, of GATA6/PDGFRA. (h) Heatmap of endodermal lineages arranged by pseudotime. (i) Line plots of endoderm cells arranged by pseudotime show up-regulation of *SOX17*/*FOXA2* with concurrent downregulation of *POU5F1*.

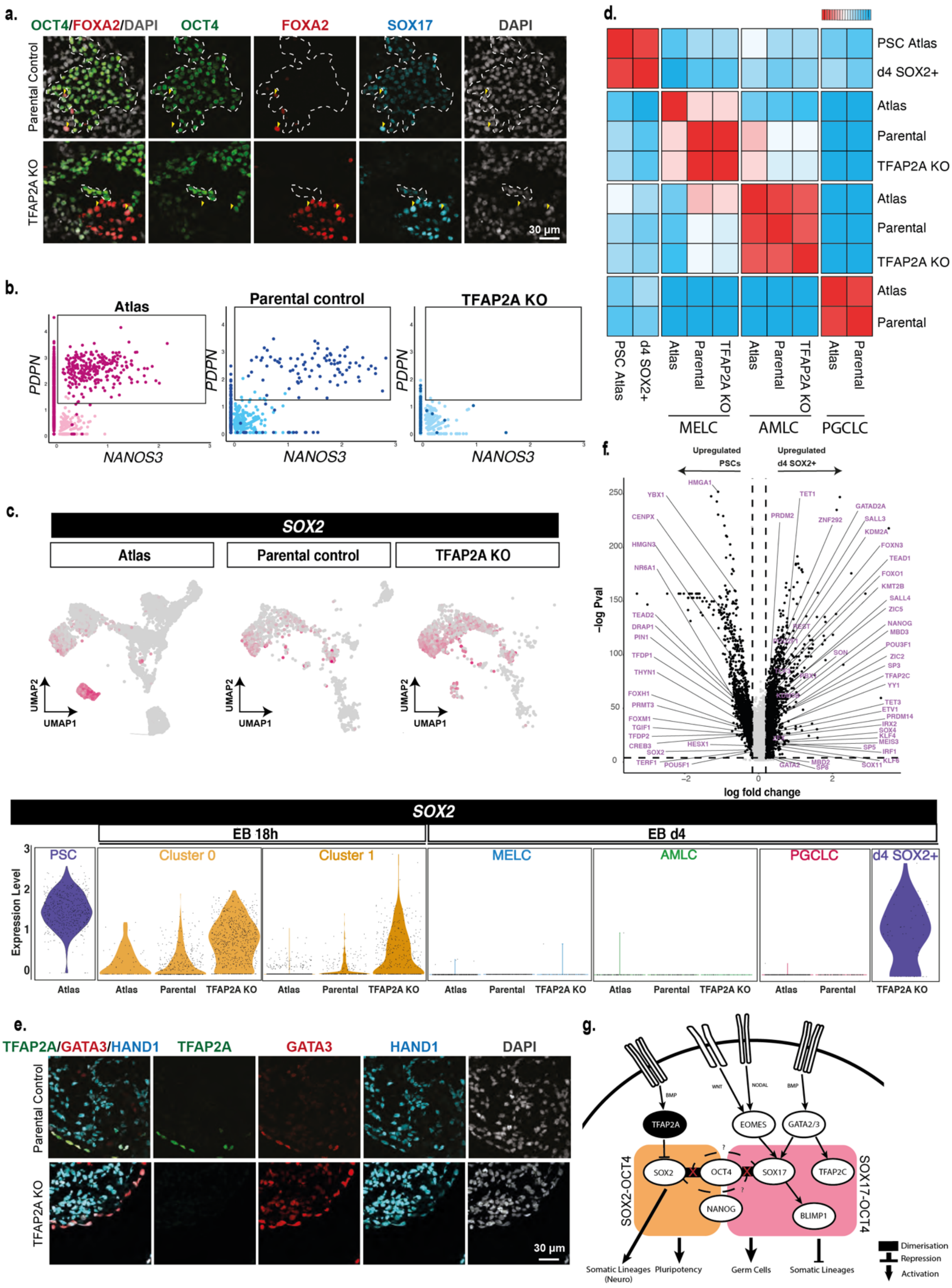


**Supplementary Figure 7:**
(a) Immunofluorescence shows PGCLC (SOX17/OCT4+), endoderm and (SOX17/FOXA2+) in TFAP2A KO d4 embryoid bodies. (b) Scatter plots of *NANOS3* versus *PDPN* levels in the single-cell data shows a lack of double-positive cells, indicating an absence of PGCLCs in the TFAP2A KO. (c) UMAP and accompanying violin plot showing SOX2 expression for the reference, parental control, and TFAP2A KO lines. (d) Pseudobulk cross-correlation heatmap of individual cell lineages between the reference dataset, parental line, and TFAP2A KO lines show consistent behaviour in specified MELC and AMLC lineages across all three lines, with PGCLCs showing consistent behaviour between the reference line and parental line. (e) Immunofluorescence shows the presence of amnion (GATA3+) and mesoderm cells (HAND1+) in TFAP2A KO d4 embryoid bodies. (f) Volcano plot for differentially expressed genes between the d4 SOX2+ cluster in TFAP2A KO vs PSCs (in parental control). (g) A model of the transcription factor network necessary for human PGCLC specification.

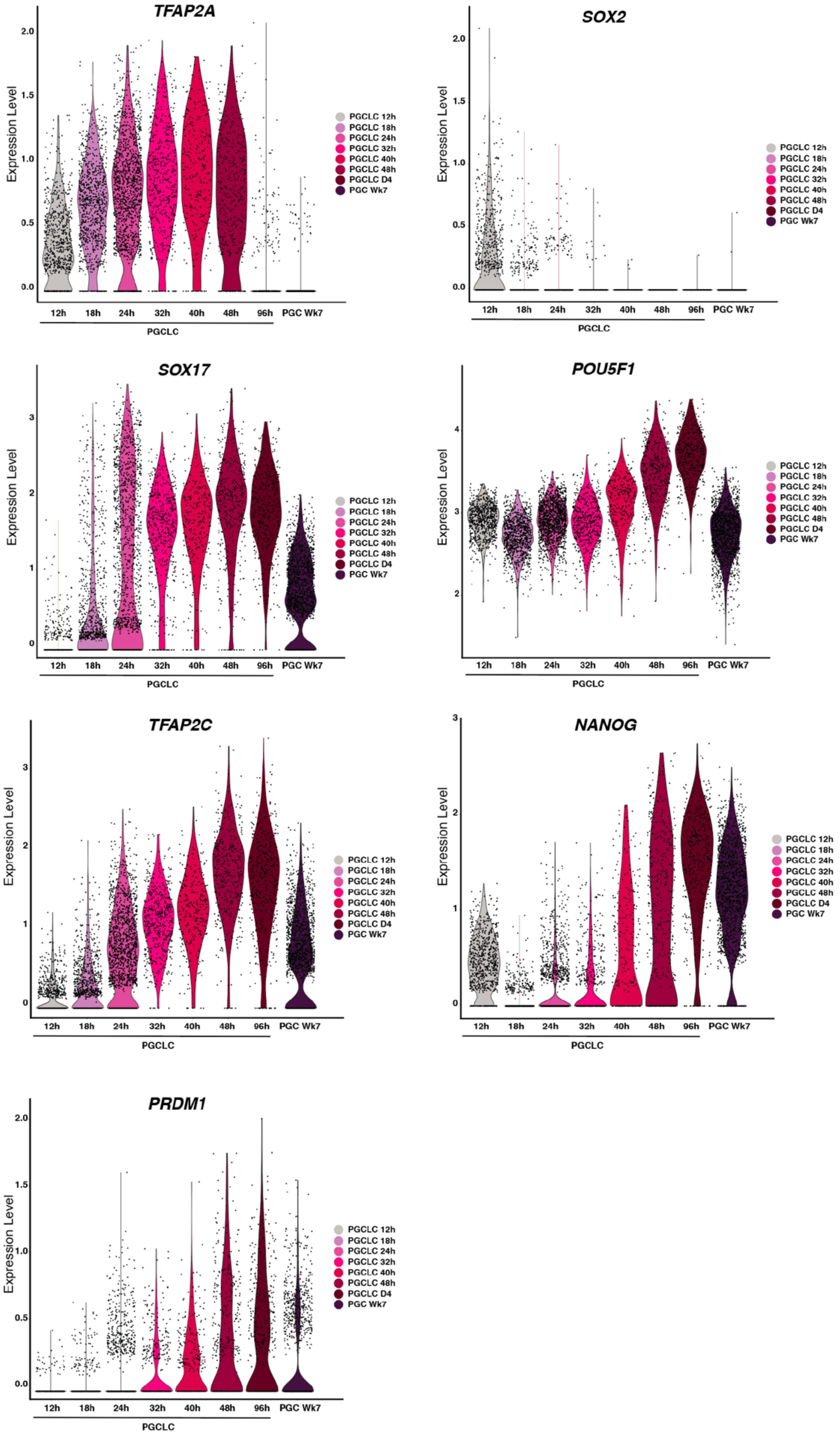


**Supplementary Figure 8:**
Violin plots for TF expression in the PGCLC trajectory.

**Supplementary Table 1: Datasets used for the gastrulation and germ cell roadmap**

| Dataset | Paper | Type | Sample type | Cell types |
|---|---|---|---|---|
| **Competent populations** | Our dataset | 10X | Competent populations | PreME, ME |
| **Embryoid bodies** | Our dataset | 10X | Embryoid bodies | PSLC,MELC, AMLC, PGCLC, 4i, PSC, PGC, DELC |
| **Embryoid bodies** | Our dataset | 10X | Embryoid bodies 18h and 48h (TFAP2A KO line and parental cell line) | PSLC,MELC, AMLC, PGCLC |
| **Embryoid bodies** | Chen et al. (2019) | 10X | Embryoid bodies, competent populations in low competence cell lines | PSLC, MELC, AMLC, PGCLC, PSC, DELC |
| **Microfluidic amnioids** | Zheng et al. (2019) | 10X | Microfluidic amnioids and PSCs | PSLC, MELC, AMLC, PGCLC, PSC, DELC |
| **2D gastruloids** | Minn et al. (2019) | 10X | Micropatterned gastruloids (48h) | PSLC, MELC, AMLC, PGCLC, PSC, DELC |
| **Human embryos** | Vento-Tormo et al. (2018) | 10X | Trophoblast | Tb |
| **Human embryos** | Xiang et al. (2019) | Smart-seq2 | *In vitro* human embryo culture | Epiblast, primitive streak, amnion |
| **Human embryos** | Zhou et al. (2019) | Smart-seq2 | *In vitro* human embryo culture | Epiblast, primitive streak |
| **Cynomolgus embryos** | Ma et al. (2019) | Smart-seq2 | *In vitro* cynomolgus embryo culture | Epiblast, primitive streak, PGCs, amnion |
| **Cynomolgus embryos** | Nakamura et al. (2016) | Smart-seq2 | *In vivo* cynomlgus embryos | Epiblast, primitive streak, PGCs |
| **Human gonadal samples** | Guo et al. (2015) | Smart-seq2 | Human gonadal PGCs | PGCs |
| **Human gonadal samples** | Li et al. (2019) | Smart-seq2 | Human gonadal PGCs | PGCs |
| **Human gastrula** | Tyser et al. (2020) | Smart-seq2 | Human CS7 gastrula | Epiblast, primitive streak, mesoderm, endoderm, amnion, PGCs |
| **Marmoset gastrula** | Bergmann et al. (2020) | Smart-seq2 | Marmoset CS5, CS6, CS7 gastrula | Epiblast, primitive streak, mesoderm, endoderm, amnion, PGCs |
